# A Systematic Approach to the Discovery of Protein-Protein Interaction Stabilizers

**DOI:** 10.1101/2023.01.29.526112

**Authors:** Dyana N. Kenanova, Emira J. Visser, Johanna M. Virta, Eline Sijbesma, Federica Centorrino, Holly R. Vickery, Mengqi Zhong, R. Jeffrey Neitz, Luc Brunsveld, Christian Ottmann, Michelle R. Arkin

## Abstract

Protein-protein interactions (PPIs) are responsible for the proper function of biological processes and, when dysregulated, commonly lead to disease. PPI stabilization has only recently been systematically explored for drug discovery despite being a powerful approach to selectively target intrinsically disordered proteins and hub proteins, like 14-3-3, with multiple interaction partners. Disulfide tethering is a site-directed fragment-based drug discovery (FBDD) methodology for screening small molecules in a quantitative, high-throughput manner. We explore the scope of the disulfide tethering technology for the discovery of selective fragments as starting points for the development of potent small molecule PPI stabilizers and molecular glues using the hub protein 14-3-3σ. The complexes with 5 biologically and structurally diverse phospho-peptides, derived from the 14-3-3 client proteins ERα, FOXO1, C-RAF, USP8, and SOS1, were screened for hit identification. Stabilizing fragments could be found for 4/5 client complexes with a diversified hit-rate and stabilizing efficacy for the different 14-3-3/client phospho-peptides. Extensive structural elucidation revealed the ability and adaptivity of the peptide to make productive interactions with the tethered fragments as key criterion for cooperative complex formation. We validated eight fragment stabilizers, six of which showed selectivity for one phospho-peptide client, and structurally characterized two nonselective hits and four fragments that selectively stabilized C-RAF or FOXO1. The most efficacious of these fragments increased 14-3-3σ/C-RAF phospho-peptide affinity by 430-fold. Disulfide tethering to the wildtype C38 in 14-3-3σ provided diverse structures for future optimization of 14-3-3/client stabilizers and highlighted a systematic method to discover molecular glues.

## INTRODUCTION

Protein-protein interactions (PPIs) are essential to biology and their dysregulation is central to many diseases including cancer and neurodegeneration.^1–4^ Many of these important PPIs include “hub proteins” that interact with a large number of protein partners, ranging from a few dozen to a few thousand.^5^ Small molecules that inhibit or stabilize individual PPIs within these networks would be powerful tools to understand the effect of a single PPI on cellular function. Although PPIs were historically considered “undruggable”, there has been much progress in developing small molecule PPI inhibitors as biological probes and therapeutics.^6–10^ By contrast, PPI stabilization has remained largely under-explored, despite its potential to be a selective method for the manipulation of a single interaction within a protein network.^11,12^ Stabilization also has the potential to target unstructured, difficult to drug proteins via composite PPI binding pockets.^13,14^ Molecular glue degraders and natural products have demonstrated the therapeutic value of stabilizing native or non-native (neomorphic) PPIs.^15–17^ However, there are few robust, generalizable strategies to discover PPI stabilizers prospectively.^11,18^ Here, we describe a robust and instructive approach, using site-directed fragment based drug discovery (FBDD) to systematically discover molecular glues.

FBDD is a well-established method for the discovery of small molecules towards challenging targets.^19,20^ Fragments are simple chemical building blocks that – owing to their small number of atoms – sample chemical space efficiently. FBDD involves screening for weakly binding fragments that target subsites within a binding site, followed by fragment optimization via linking two fragments or elaborating a fragment-sized core. Disulfide tethering is a method of FBDD that capitalizes on a native or engineered cysteine residue proximal to an envisaged ligand binding site.^21–24^ In the context of orthosteric PPI stabilization, this binding site is composed of both members of the protein complex (the composite PPI interface). Fragments that bind to this site with the correct positioning to form a protein-fragment disulfide bond are detected by intact protein mass spectrometry (MS) in a high-throughput screen.^25^ We utilize a library of approximately 1600 disulfide molecules with diverse fragments and linkers between the fragment and the disulfide.^26^ To test the efficacy of this technology to discover PPI stabilizers, we have selected the hub protein 14-3-3 and a set of its diverse partner proteins.

14-3-3 is ubiquitously expressed in mammals and plays multiple roles within the cell, including phosphorylation protection, conformational changes, subcellular trafficking, and induction or disruption of other PPIs.^13,27–29^ 14-3-3 typically binds to a phosphorylated serine/threonine of intrinsically disordered regions of its clients.^30^ With several hundred known interacting partners, the 14-3-3-binding proteome provides diverse PPI interfaces with which to test the scope and limitations of our screening technology. Furthermore, 14-3-3/client stabilization could lead to therapeutics in a variety of disease fields including oncology, neurodegeneration, inflammation, and metabolic disease.^29,31^ Previous studies using natural products such as Fusicoccin (FC-A) and Cotylenin-A (CN-A) have shown that stabilizing 14-3-3/client interactions regulates the activity of important cell signaling pathways including estrogen receptor α (ERα) and C-RAF, respectively.^14,32^

We recently demonstrated the utility of disulfide tethering to identify molecular glues of the 14-3-3/ERα PPI. We discovered a series of disulfide fragments that stabilized the complex when bound to an engineered cysteine residue in the binding groove of 14-3-3, enhancing binding of the ERα C-terminal phosphopeptide up to 40-fold.^25^ We now focus on targeting the native cysteine found in the 14-3-3 sigma isoform (14-3-3σ), which offers greater translatability for covalent molecules. Of the 7 isoforms found in mammalian cells, 14-3-3σ is the only one that harbors a cysteine residue proximal to the client binding groove, providing an additional degree of isoform specificity (Figure 1A).^30^ The Protein Data Bank contains dozens of crystallographic structures of 14-3-3 with bound phospho-peptides derived from many of its binding partners, as well as a few examples of CryoEM structures of full length proteins.^13,33–35^ This wealth of structural information allows for direct visualization of the various 14-3-3/client binding interfaces which could be capitalized on for the discovery of selective fragment stabilizers and the development of potent lead compounds through structure-guided chemical optimization. For our screens, we utilized the phospho-peptide mimetics of 14-3-3 PPI partners which bind 14-3-3 in a similar fashion to the unstructured regions of the full-length proteins, but offer greater synthetic flexibility and simplified crystallography.^13,34^

**Figure 1.**
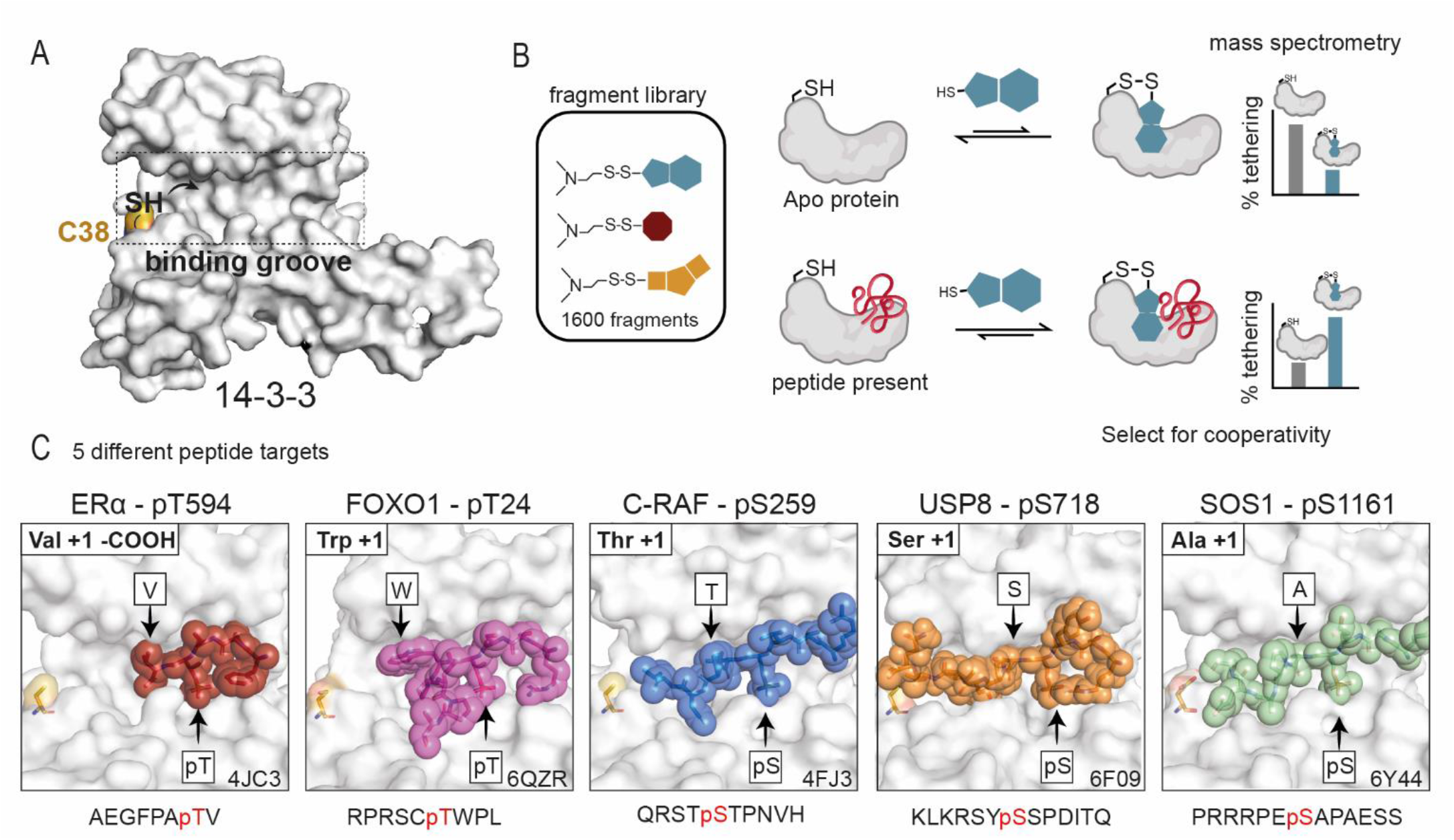
14-3-3/client stabilizer approach. (A) The client protein binding groove of a 14-3-3σ monomer (white surface) highlighting the native cysteine (C38; yellow surface) and target thiol. (B) General schematic of the primary disulfide tethering screen. Fragments were incubated with *apo* 14-3-3σ (white) without any phospho-peptide (top) and 14-3-3σ with the phospho-peptide client present (bottom). Fragments were assessed for their covalent engagement of C38 via mass spectrometry, termed “% tethering”. Fragments that bind 14-3-3σ with a higher % tethering in the presence of phospho-peptide than in the *apo* screen are selected for further analysis of cooperativity. (C) Crystallographic structures of the 5 phospho-peptide clients bound in the 14-3-3σ (white surface) binding pocket showing proximity to C38 (yellow surface). ERα (red sticks) has a C-terminal motif with phospho-threonine (pT) in the penultimate position and C-terminal valine (V) in the +1 position. FOXO1 (pink sticks) has a curved motif with tryptophan (W) in +1 position. C-RAF (blue sticks), USP8 (orange sticks), and SOS1 (green sticks) extend to various degrees into the 14-3-3 binding groove, with threonine (T), serine (S), and alanine (A) residues in the +1 position, respectively. PDB left to right: 4JC3, 6QZR, 4FJ3, 6F09, 6Y44.

Here, we used the disulfide tethering technology to systematically achieve selective PPI stabilization of 14-3-3 client phospho-peptides with diverse sequences and structures. The selected clients are also modulated by 14-3-3 in a way that could be therapeutically useful in cancer, metabolic disease, and/or rare disease.^14,36–39^ For four of the five targets, effective PPI stabilizers were identified. Crystallographic and functional data highlight the molecular recognition of fragments for the distinctive composite PPI interfaces formed by 14-3-3 bound to client phospho-peptides. In particular, the C-RAF- and FOXO1-based peptide-protein interactions with 14-3-3 yielded fragments with high selectivity and/or stabilization factors. The diversity of sequences and conformations found in 14-3-3/client complexes make the 14-3-3 interactome particularly promising for small-molecule PPI stabilization; furthermore, the disulfide tethering approach is remarkably effective at selecting chemical starting points for further design of potent and selective PPI stabilizers.

## RESULTS AND DISCUSSION

### Primary Screen for 14-3-3/Client Stabilizers

The disulfide tethering screen targeted C38, a native cysteine on 14-3-3σ located proximal to the natural product binding pocket within the phospho-peptide recognition groove (Figure 1A, Figure S1). The cysteine forms a reversible covalent bond with the fragment thiol through disulfide exchange; the amount of bound fragment is measured by MS. A fragment stabilizer is expected to show a higher “% tethering” in the presence of the 14-3-3σ/client phospho-peptide complex than 14-3-3σ alone due to cooperativity between the fragment and the peptide (Figure 1B). The screening was performed on five different peptide targets displaying three conceptually distinct 14-3-3 interaction motifs (Figure 1C): truncated (ERα),^14,40^ turned (FOXO1),^37^ and linear (C-RAF, USP8, SOS1).^32,35,38,41^

14-3-3σ (100 nM) was screened in complex with the 5 client phospho-peptides at a concentration twice their respective K_D_ values (Figure S2). This condition provided a consistent presence of the 14-3-3σ/phospho-peptide composite interface that the fragments would engage. The 14-3-3σ/phospho-peptide complex was incubated with a single concentration of fragment (200 μM) under reducing conditions (250 μM β-mercaptoethanol) for 3 hours before samples were measured by intact-protein LC/MS. The % tethering threshold for hit selection was three standard deviations (3*SD) above the average % tethering for that condition (Figure 2A). In the quadrant of highest interest, potential stabilizing fragments showed % tethering above the tethering threshold in the peptide screen and % tethering below the tethering threshold in the *apo* screen (Figure 2B, green quadrant). Neutral compounds showed significant % tethering for both 14-3-3σ/phospho-peptide and *apo* (Figure 2B, yellow quadrant). Potential inhibitory fragments showed significant % tethering above the tethering threshold in the *apo* screen but not in the presence of peptide (Figures 2B, red quadrant). Compounds were clustered in a heat map based on % tethering in each of the five peptide screens and *apo* 14-3-3 screen (Figure 2C). An overlapping fragment hit cluster was identified for ERα, USP8, and SOS1 (Figure 2C, green box), whereas a cluster of unique hit fragments was identified for both C-RAF and FOXO1 (Figure 2C, yellow boxes), indicating a difference in the abundance of selective stabilizers from the primary screens.

**Figure 2.**
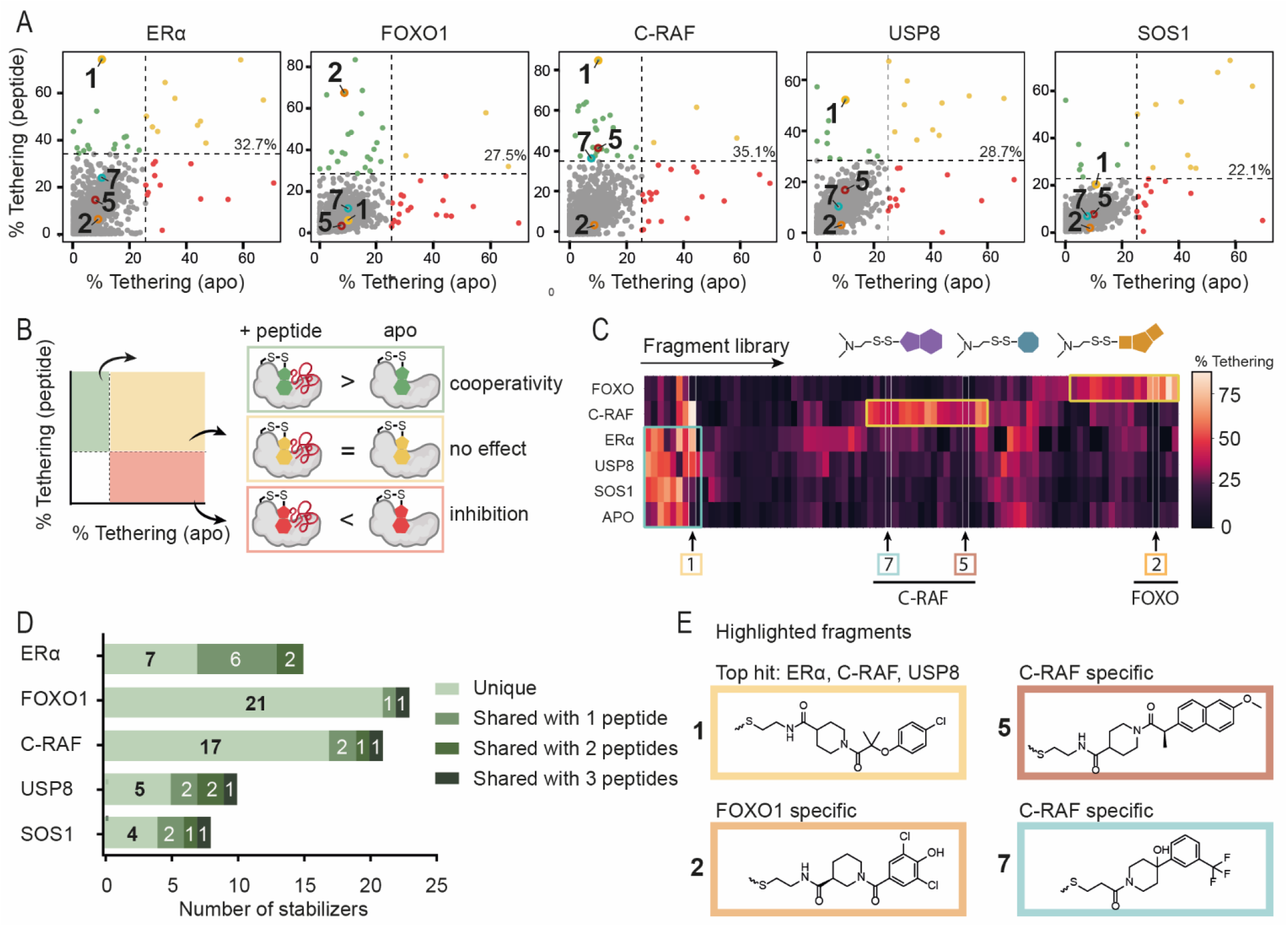
Primary tethering screen results. (A) Scatterplot data illustrating the correlation of % tethering of fragments to 14-3-3σ in the presence of the phospho-peptide (y-axis) as compared to *apo* 14-3-3σ (x-axis). Hit selection threshold (mean + 3*SD) in each screen is indicated by a black dashed line. Compounds **1, 2, 5** and **7** are indicated as yellow, orange, red and cyan circles, respectively. (B) Schematic of compound scatterplots. Quadrants are outlined by dotted lines signifying 3*SD above average % tethering for compounds in the presence of phospho-peptide (horizontal line) and *apo* 14-3-3σ (vertical line). Compounds in green quadrant showed increased binding to 14-3-3σ in the presence of phospho-peptide, yellow quadrant showed neutral binding to 14-3-3σ, and red quadrant showed a reduced binding in presence of phospho-peptide. (C) Heat map of hit fragments across all 5 phospho-peptide screens and *apo* 14-3-3σ screen. Compounds clustered based on % tethering in each screen. Compounds **1, 2, 5** and **7** were of primary interest as non-selective and selective stabilizers. (D) Number of stabilizers of each peptide that were: unique, shared with one other peptide, shared with two other peptides, or shared with three other peptides (green bars with the darker color shared with more peptides). (E) Chemical structures of highlighted fragment hits **1, 2, 5**, and **7**.

Each 14-3-3σ/phospho-peptide screen yielded potential stabilizing fragments, but the number and binding efficiency varied (Figure 2A, 2C, 2D, and Tables S1-S5). The initial screen for ERα yielded 15 hit fragments including 7 unique stabilizers and a 33% tethering threshold. The FOXO1 screen yielded 23 hit fragments including 21 unique stabilizers and a 28% tethering threshold. The C-RAF screen yielded 21 fragments including 16 unique stabilizers and a 35% tethering threshold. The USP8 screen yielded 10 hit fragments including 5 unique stabilizers and a 29% tethering threshold. The SOS1 screen yielded 8 hit fragments including 4 unique stabilizers and a 22% tethering threshold (Figures 2A and 2D). Figure 2E depicts representative chemical structures for each target.

### Non-Selective Stabilizing Compound 1

In the initial screen, compound **1** was identified as top hit for ERα, C-RAF and USP8 (Figure 2). **1** was further characterized by three dose-response experiments. Mass spectrometry (MSDR, analyzing fragment binding to protein, quantified by DR_50_ values) and fluorescent anisotropy (FADR, analyzing peptide binding to protein in the presence of compound, quantified by EC_50_ values) defined the binding affinity for the fragment and its effective concentration, respectively (Figure 3B). The compound’s effect on the 14-3-3/client PPI was then determined by titrating 14-3-3 in a fluorescence anisotropy assay at constant peptide and compound concentrations (quantified by K_D_app_). In all three validation assays, **1** displayed a strong preference for C-RAF, followed by ERα and USP8, and had no activity with FOXO1 or SOS1. Compound **1** showed DR_50_ values of 7 nM for C-RAF, 18.1 μM for ERα, and 24 nM for USP8 (Figure 3C) as well as EC_50_ values of 922 nM for C-RAF, 1.31 μM for ERα, and 3.38μM for USP8 (Figure S3). In the protein titrations, **1** increased peptide affinity for 14-3-3σ by 81-fold in the C-RAF complex, 19-fold for 14-3-3σ/ERα, and 4-fold for 14-3-3σ/USP8 (Figure 3D and Table 1).

**Table 1.** Tethering and stabilization of 14-3-3σ/clients by compound 1.

| Peptide | MSDR<br>(250 μM<br>BME) | FADR<br>(50 μM<br>BME) | Protein Titrations<br>(50 μM BME) |  | Fold<br>Stab. |
| --- | --- | --- | --- | --- | --- |
|  | DR<sub>50</sub><br>(μM) | EC<sub>50</sub><br>(μM) | K<sub>D_app</sub> | K<sub>D_DMSO</sub> |  |
| CRAF | 0.007 | 0.922 | 106 nM | 8.5 μM | 81 |
| ERα | 18.1 | 1.31 | 21 nM | 360 nM | 19 |
| USP8 | 0.024 | 3.38 | 1.1 μM | 4.5 μM | 4 |
| SOS1 | >2 mM | >2 mM | N/A | N/A | N/A |
| FOXO1 | >2 mM | >2 mM | N/A | N/A | N/A |
| apo | >2 mM | >2 mM | N/A | N/A | N/A |

**Figure 3:**
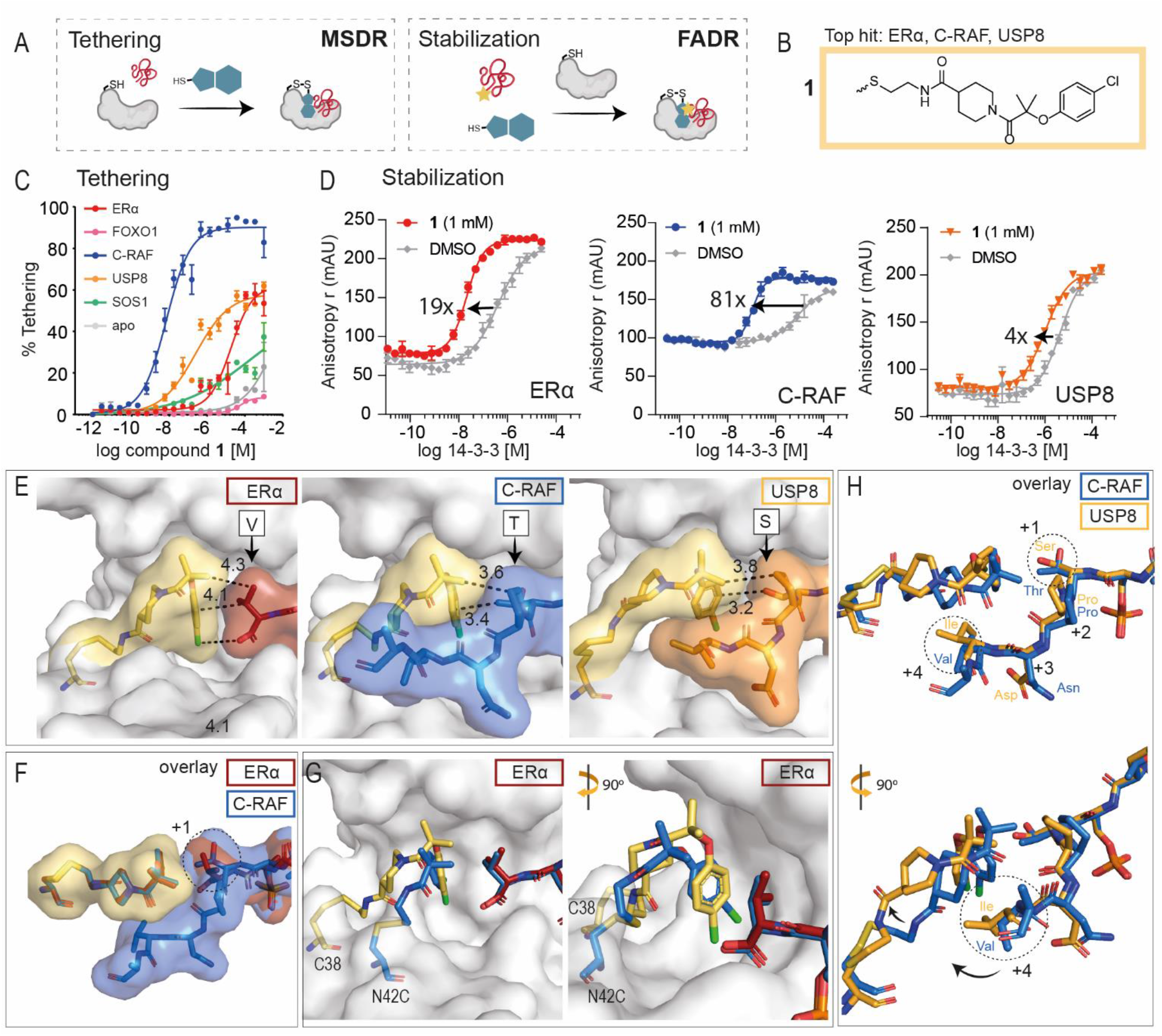
Overview of biochemical and structural properties of non-selective stabilizer **1**. (A) In mass spec dose response (MSDR), the focus was on compound binding to 14-3-3σ, measured by % tethering; fluorescence anisotropy dose response (FADR) experiments determined degree of stabilization, measured by phospho-peptide binding to 14-3-3σ in presence of compound. (B) Chemical structure of stabilizer **1**. (C) MSDR curves for **1** showing percentage of fragment/protein conjugate formation with 14-3-3σ *apo*, or in the presence of ERα, FOXO1, C-RAF, USP8 or SOS1 peptide. (D) 14-3-3σ titrations to fluorescein-labeled ERα, C-RAF or USP8 in the presence of DMSO or **1** (1 mM), reporting a 19-, 81-, and 4-fold increase of the 14-3-3σ/peptide binding interface, respectively. (E) Crystal structure of **1** bound to 14-3-3σ C38 in complex with (from left to right) ERα peptide, C-RAF peptide, and USP8 peptide. Distances are indicated (Å, black dashes). (F) Overlay of **1’s** conformations when interacting with ERα and C-RAF. (G) Overlay of **1** (yellow) bound to 14-3-3σ C38 and previously reported stabilizer (blue) bound to 14-3-3σ mutant N42C (PDB ID: 6HMT) interacting with ERα phospho-peptides. (H) Overlay of **1** bound to 14-3-3σ C38 interacting with C-RAF and USP8.

Crystal structures for compound **1** were obtained by co-crystallizing with ERα, C-RAF, or USP8 bound to 14-3-3σ (Figure 3E), with clear density for both **1** and the peptides (Figure S4). Comparing the three co-crystal structures, the strongest electron density and ligand occupancy for **1** was observed in the co-crystal structure with ERα. For ERα, the phenyl ring of **1** stacked against the +1 Val with a distance of ∼4 Å (Figure 3E). Compound **1** showed an identical binding mode in the presence of C-RAF (Figure 3F), for which the +1 Thr was 3.5 Å from the phenyl ring, while the remainder of the C-RAF peptide wrapped around the fragment. These additional hydrophobic interactions could explain the higher fold stabilization with the C-RAF peptide compared to ERα (Figure 3D, 3F). Interestingly, **1** shared the binding moiety with N42C-tethered stabilizers that were discovered previously for ERα (Figure 3G).^25^ Whereas compound **1’**s chloro-group was not positioned identically, the longer linker of **1** bridged the larger distance from C38 compared to N42C. In the presence of the USP8 peptide, the phenyl ring of **1** was turned, thereby shifting the fragment up and back into the 14-3-3σ pocket (Figure 3E). This conformational change seemed necessary because the USP8 peptide allowed for less space (Figure 3H). While the +1 Ser of UPS8 did not show any specific interaction with **1**, its +4 Ile pushed the fragment towards 14-3-3σ, which was not an ideal position for this fragment as was reflected by the weak electron density and the minimal stabilization for USP8. By contrast, the +4 Val of C-RAF allowed for more space, thereby positioning **1** in a preferred conformation. It is noteworthy that **1** did not stabilize FOXO1 or SOS1 to 14-3-3σ. A crystallographic overlay of **1** with the FOXO1 peptide showed a steric clash with the +1 Trp of FOXO1, explaining its lack of stabilization (Figure S5A). In contrast, the +1 Ala residue of SOS1 would not contact the phenyl ring of **1**, perhaps explaining why no stabilization was observed (Figure S5B).

### FOXO1 Selective Stabilizers

The FOXO1 peptide showed the highest number of stabilizing hits in our initial screen. For FOXO1, of the 23 initial stabilizers, 21 showed selectivity for the 14-3-3σ/FOXO1 phospho-peptide complex over *apo* 14-3-3σ and the other phospho-peptide clients in the initial screen (Figure 2D). Interestingly, the unique 21 FOXO-stabilizers had a highly conserved scaffold, with the phenyl ring engaging FOXO1 often decorated with halogens or a triazole moiety (Figure S6). Eight of these compounds were validated in the MSDR (Figure S7). Of the eight compounds, five compounds had enough material to retest and were active in the FADR assays (Figure 4A, Figure S8, and Table S2). The binding affinity of compound **2** to 14-3-3σ was >10,000-fold better in the presence of the FOXO1 phospho-peptide than *apo* 14-3-3σ and all other phospho-peptide clients (DR_50_ = 360 nM vs. >2 mM; Figure 4B and Table 2). Compounds **3** and **4** had DR_50_ values >450-fold and >2,000-fold better, respectively (Table 2 and Figure S7). Compounds **2, 3**, and **4** showed the greatest fold-stabilization in the protein titrations decreasing 14-3-3σ/FOXO1 K_D_ values 5-fold, 4-fold, and 12-fold (Figure 4C, Table 2, and Figure S9). It should be noted that while a high % tethering was observed for the FOXO1 stabilizers, the protein titrations only showed a modest shift in stabilization. This is likely due to the tight binding of the FOXO1 phospho-peptide, with a K_D_ value of 50 nM, already close to the limit of detection of this assay.

**Table 2.** Properties of selective FOXO1 stabilizers.

| Cmpnd | MSDR<br>(250 μM<br>BME) | FADR<br>(50 μM<br>BME) | Protein Titrations<br>(50 μM BME) |  |  |
| --- | --- | --- | --- | --- | --- |
|  | DR<sub>50</sub><br>(μM) | EC<sub>50</sub><br>(μM) | K<sub>D,app</sub> | K<sub>D,DMSO</sub>* | Fold Stab. |
| 2 | 0.36 | 5.10 | 7.7 nM | 39 nM | 5 |
| 3 | 2.2 | N/A | 10.6 nM | 42 nM | 4 |
| 4 | 143 | N/A | 9.7 nM | 111 nM | 12 |

\*K<sub>D</sub> for peptide is accurate within 3-fold range. These values are shown on the same plate as protein titrations with compound.

**Figure 4:**
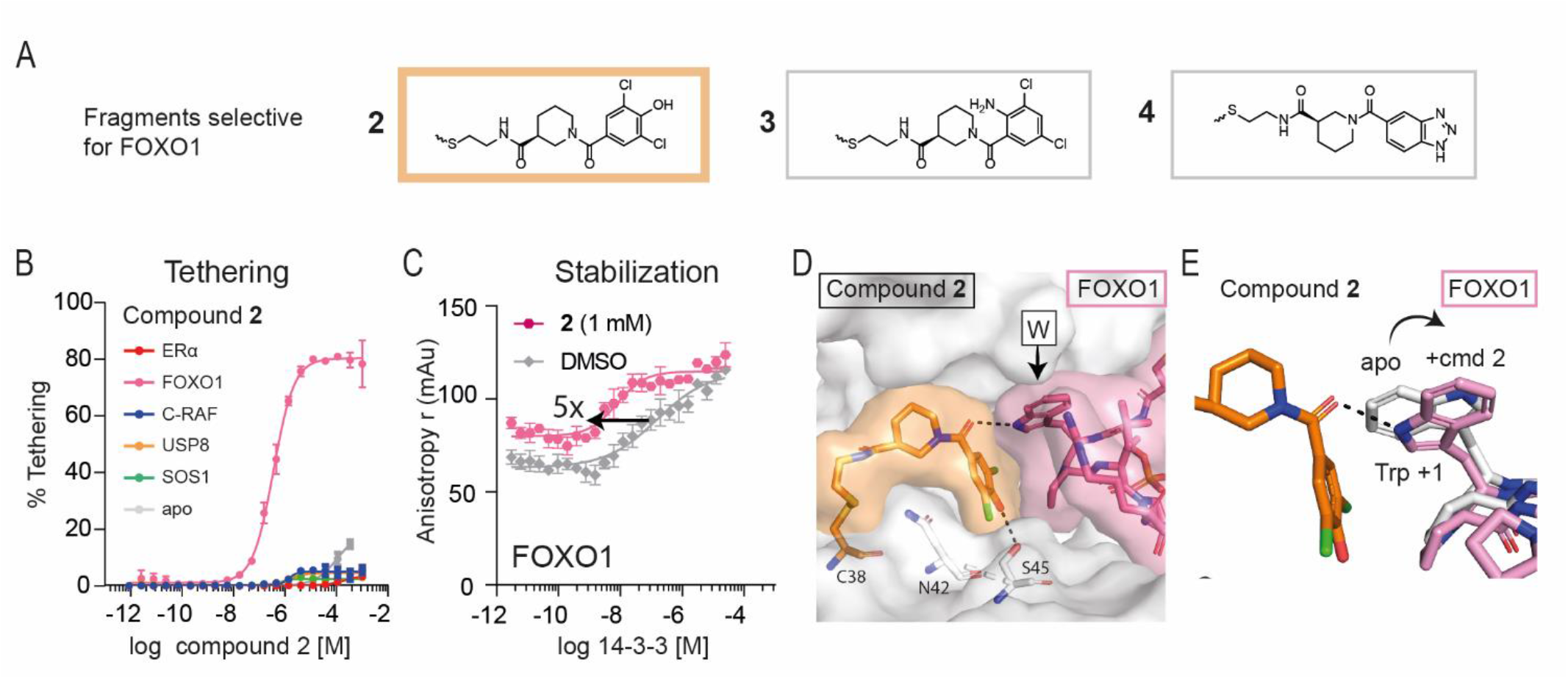
Overview of selective stabilizers for FOXO1. (A) Chemical structures of highlighted FOXO1 selective stabilizers **2-4**. (B) MSDR curves for **2** showing percentage of fragment/protein conjugate formation with 14-3-3σ *apo*, or in the presence of ERα, FOXO1, C-RAF, USP8 or SOS1 peptide. (C) 14-3-3σ titrations to fluorescein-labeled FOXO1 in the presence of DMSO or **2** (1 mM), reporting a 5-fold increase in 14-3-3σ/FOXO1 binding. (D) Crystal structure of **2** (orange) bound to 14-3-3σ (white) C38 in complex with FOXO1 phosphopeptide (pink). (E) Overlay of FOXO1 peptide in the apo-structure (white) with the FOXO peptide (pink) in presence of **2** (orange).

A co-crystal structure for FOXO1/**2**/14-3-3σ was obtained, with clear density for both **2** and the FOXO1 peptide (Figure S10A). The phenyl ring of **2** stacked against the front of the FOXO1 peptide consisting of the +1 Trp and the +2 Pro residues (Figure 4D). Strikingly, in the presence of **2**, the Trp of FOXO1 underwent a conformational change to form a hydrogen bond with its NH and the amide carbonyl of **2** (Figure 4E). Moreover, the hydroxyl on the phenyl ring of **2** made a hydrogen bond with the S45 of 14-3-3σ, explaining the benefit of a hydrogen donor or, potentially, acceptor at that position. Compound **3** was also co-crystallized with FOXO1 (Figure S10B), showing a highly similar binding mode, but a lack of the hydrogen bonding with S45 of 14-3-3σ (Figure S10C). An overlay of **2** with the other peptides revealed that **2** could not reach the smaller +1 residues in the other client peptides or that the peptides sterically clashed (Figure S11), potentially explaining its selectivity for FOXO1 over the other peptides. Previous work discovered imine-based stabilizers for the 14-3-3/Pin-1 complex which, similar to FOXO1, has a +1 Trp.^42^ In that work, the Trp engaged in π-π stacking interactions with an aromatic ring of the stabilizers. By contrast, the +2 Pro of FOXO1 locked the conformation of the +1 Trp and thereby prevented such a π-π stacking interaction with **2**, while the +2 Arg of Pin-1 allowed π-π stacking to take place. Thus, while the compound **2**/**3** scaffold emphasized the chemical moieties necessary for stabilizing FOXO1, crystal structures also expose a lack of flexibility of the FOXO1 peptide.

### C-RAF Selective Stabilizers

Following FOXO1, C-RAF had the highest number of stabilizers. Of the 21 initial C-RAF stabilizers, 16 compounds showed selectivity for the 14-3-3σ/C-RAF phospho-peptide complex over *apo* 14-3-3σ and the other phospho-peptide clients in the primary screen (Figure 2D). Eleven compounds showed a similar scaffold which was remarkably analogous to the conserved scaffold for the FOXO1 stabilizers (Figure S12). However, the linker element of these compounds was often longer in the case for C-RAF, and the phenyl ring was decorated with large cyclic groups while for FOXO1 only smaller halogen groups were tolerated. This is likely due to the smaller +1 residue of C-RAF (Thr for C-RAF, Trp for FOXO1), thereby leaving more space for the compound. Furthermore, two C-RAF stabilizers were shared with ERα, both of which have a similar size in +1 residue (Val for ERα, Thr for C-RAF). Nine of the 16 selective compounds were validated for potency and selectivity in the MSDR (Figure S13). Four of the nine compounds (compounds **5**-**8**; Figure 5A) showed activity in FADR (Figure S14 and Table S3) and stabilization in the protein titrations (Table 3 and Figure S15).

**Table 3.**
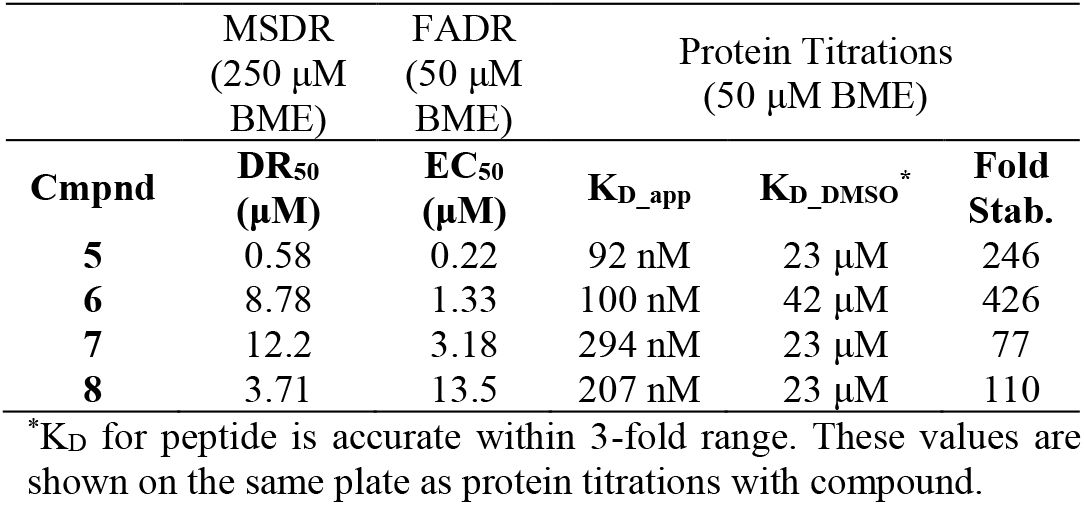
Properties of selective C-RAF stabilizers.

**Figure 5:**
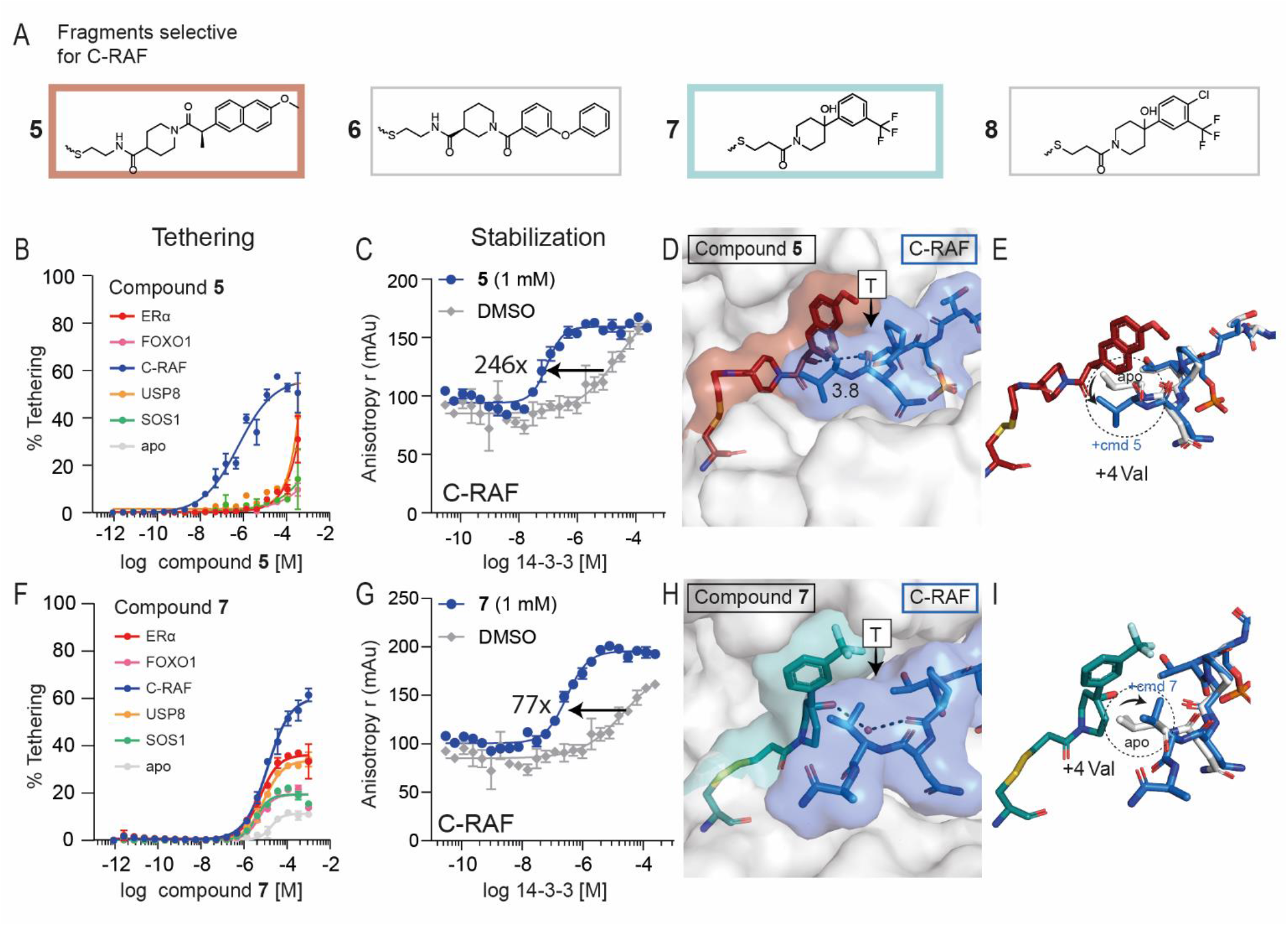
Overview of selective stabilizers for C-RAF. (A) Chemical structures of highlighted C-RAF selective stabilizers **5-8**. (B) MSDR curves for **5** showing percentage of fragment/protein conjugate formation with 14-3-3σ *apo*, or in the presence of ERα, FOXO1, C-RAF, USP8 or SOS1 peptide. (C) 14-3-3σ titration to fluorescein-labeled C-RAF in the presence of DMSO or **5** (1 mM), reporting a 246-fold increase of 14-3-3σ/C-RAF binding. (D) Crystal structure of **5** (red) bound to 14-3-3σ (white) in complex with C-RAF phospho-peptide (blue). (E) Overlay of C-RAF peptide in the apo-structure (white) with the C-RAF peptide (blue) in presence of **5** (red). (F) MSDR curves for **7** showing percentage of fragment/protein conjugate formation with 14-3-3σ *apo*, or in the presence of ERα, FOXO1, C-RAF, USP8 or SOS1 peptide. (G) 14-3-3σ titration to fluorescein-labeled C-RAF in the presence of DMSO or **7** (1 mM), reporting a 77-fold increase of 14-3-3σ/C-RAF binding (H) Crystal structure of **7** (teal) bound to 14-3-3σ (white) in complex with C-RAF phospho-peptide (blue). (I) Overlay of C-RAF peptide in the apo-structure (white) with the C-RAF peptide (blue) in presence of **7** (teal).

Compounds **5** and **6** were the most effective stabilizers. Compound **5** had a DR_50_ value >3,000-fold lower in the presence of the C-RAF peptide compared to 14-3-3σ alone (Figure 5B) and showed a 246-fold stabilization of the 14-3-3σ/C-RAF phospho-peptide complex (K_D_ = 23 μm to 92 nM; Figure 5C). Compound **6** had a DR_50_ value 230-fold lower in the presence of C-RAF compared to *apo* 14-3-3σ and a 426-fold stabilization of the 14-3-3σ/C-RAF complex (Table 3, Figure S13 and S15).

The crystal structure of **5** with C-RAF and 14-3-3σ revealed a contact between the naphthalene ring of **5** and the +1 Thr residue of C-RAF. The methyl group of **5** also seems important for hydrophic interactions with the +1 Thr residue of C-RAF, at a distance of 3.8 Å (Figure 5D, Figure S16A). An overlay of the C-RAF peptide in the presence of **5** with the *apo* C-RAF peptide showed no change in conformation of the +1 Thr residue. In contrast, the +4 Val residue of the C-RAF peptide changed conformation to make space for **5** (Figure 5E).

We also crystallized compound **7** as a representative of the other structural class of the selective C-RAF stabilizers (Figure S16B). Compound **7** had a DR_50_ value >228-fold lower in the presence of the C-RAF peptide than apo 14-3-3σ (Figure 5F) and was less selective for C-RAF compared to **5** in the MSDR (Figure S13). However, compound **7** showed no stabilization of any of the peptides other than C-RAF in the FADR (Figure S14C), reflecting the selectivity shown in the primary screen. The weaker 14-3-3σ binding of **7** (12.2 μM DR_50_) was reflected in a somewhat lower stabilization of the 14-3-3σ/C-RAF complex compared to the other chemotype of **5** and **6** (77-fold vs 246- and 426-fold, respectively; Figure 5G, Figure S15). Co-crystallization of **7** with C-RAF and 14-3-3σ revealed a novel orientation of its phenyl ring towards the roof of 14-3-3σ, positioning its trifluoromethyl group above the C-RAF peptide (Figure 5H). While the conformations of **5** and **7** were quite different, an overlay of the two structures shows that the trifluoromethyl group of **7** occupied the same cavity as the naphthalene ring of **5** (Figure S16C). Furthermore, an overlay of the C-RAF peptide in the presence of **7** with the *apo* C-RAF peptide revealed a conformational change of the +4 Val of C-RAF, which stacked against the compound, pushing it towards 14-3-3σ. Additionally, a water-mediated hydrogen bond was formed between **7** and the backbone of C-RAF peptide (Figure 5I). The lower specificity for C-RAF of **7** in the MSDR could be due to its small size, leaving room for alternative +1 residues to have a cooperative effect on 14-3-3σ engagement. Stabilizer **8** had an almost identical structure to **7**, differing only in a chloro-group in the para-position of the phenyl ring, and showed similar binding modes to **7** in its structure with C-RAF (Figure S16D and E).

Next to these C-RAF selective stabilizers, the non-selective stabilizer compound **1** also showed a large fold-stabilization towards the C-RAF peptide (Figure 3). A crystallographic overlay of these three scaffolds revealed remarkable differences in conformation of the C-RAF/compound interactions (Figure S16F). These changes highlight the flexibility of the C-RAF peptide, perhaps leading to its facility for stabilization, especially in the case of the stabilizers’ phenyl ring, which can occupy a wide range of positions and conformations in combination with the C-RAF client phospho-peptide.

## CONCLUSIONS

Systematic methods to discover small-molecule stabilizers of PPI would enable chemical biologists to probe challenging biological systems with potency and precision. By trapping proteins in complexes, stabilization can target proteins with intrinsically disordered regions and allow manipulation of a specific PPI from among related hub protein complexes within a network. Disulfide tethering, a powerful FBDD technique, is readily tunable to a specific site on a protein of interest, amenable to HTS, and provides a direct quantitative measurement of fragment binding.

Here, we explored the scope of the disulfide tethering technology using the hub protein 14-3-3σ and 5 biologically and structurally diverse phospho-peptides derived from the 14-3-3 client proteins ERα, FOXO1, C-RAF, USP8, and SOS1. Of the 1600 fragments in the disulfide library, 62 showed activity as stabilizers for one or more phospho-peptides and were assessed by MSDR. 36 of the 62 compounds were taken forward into the FADR assays to determine stabilization of a 14-3-3 client phospho-peptide. Finally, eight compounds showed cooperativity with the 14-3-3σ/phospho-peptide complex via 14-3-3σ protein titrations, and six were structurally characterized for their contacts with 14-3-3σ and the client phospho-peptide via x-ray crystallography (Figure S17A). Thus, the disulfide tethering strategy systematically discovered stabilizers for a range of peptide sequences, conformations, and affinities.

Of the 5 peptide targets selected, we discovered stabilizers for four clients, two of which also had selective stabilizers. Fragments increased binding affinity of the 14-3-3σ/phosphopeptide complex as much as 430-fold in the case of **6** and 250-fold for our best structurally characterized hit, **5**. Selective stabilizers distinguished between phospho-peptide clients due to the unique composite binding surface created by the phosphopeptide/14-3-3σ interface (Figures 4 and 5). The non-selective stabilizers also showed varying degrees of efficacy in stabilizing different clients. Compound **1** facilitated a greater than 80-fold shift in affinity for C-RAF, a 19-fold shift for ERα, a more modest 4-fold shift for USP8, but had no effect on SOS1 and FOXO1 (Figure 3).

The individual phospho-peptide binding motifs and C-terminal residues following the phosphorylation site create a distinct environment around the 14-3-3σ C38 fragment binding pocket, dictating what chemical moieties effectively facilitated cooperativity between 14-3-3σ, the phospho-peptide client, and the fragments. The stabilizers for FOXO1 had a highly conserved scaffold, consistent with the rigidity of this peptide (Figure S17B). In contrast, the stabilizers of C-RAF were larger and showed more chemical diversity in their scaffold, emphasizing the flexibility of the C-RAF peptide. The short ERα peptide resulted in limited selectivity, sharing many stabilizers with C-RAF. Lastly, USP8 and SOS1 were the hardest to target, likely due to the proximity of the peptide C-terminus to C38 of 14-3-3σ, which was also reflected in the small scaffold of the discovered stabilizers from the primary screen (Figure S17B). Alternative cysteine tethering mutations could sample different sub-pockets to stabilize peptides which occupy more of the 14-3-3 binding groove. Taken together, the intrinsic diversity of the 14-3-3/phospho-peptide composite binding interface allowed for selectivity and precision when targeting a specific 14-3-3/client PPI.

While the focus of the screen was the discovery of fragment stabilizers, the screen also identified selective inhibitors, non-selective inhibitors, and neutral compounds for each client peptide and 14-3-3 (Figure S18). Therefore, disulfide tethering is a versatile tool that can be expanded to meet a wide range of conditions and results in hits that disrupt or stabilize PPIs. 14-3-3 provides an exciting proof of concept due to its large roster of clients, involvement in many biological processes, therapeutic potential, and extensive structural data, but the applicability of FBDD reaches beyond targeting a singular protein. It is due to this ease of access and applicability that disulfide tethering lends itself to the discovery of biological probes for PPIs and novel therapeutics for previously inaccessible biological challenges and diseases related to intrinsically disordered proteins.

## ASSOCIATED CONTENT

Supporting Information. Experimental methods, tethering and stabilization data of primary screen (Table S1 – S5), XRD data collection and refinement statistics (Table S6, S7), MSDR and FADR binding curves of validation approach (S2, S3, S7, S8, S9, S13 – S15), X-ray density and crystallographic overlay of disulfide fragments (S4, S5, S10, S11, S16), chemical structures and information of discovered fragments of primary screen (S6, S12, S17, S18, S19) (PDF). This material is available free of charge via the Internet at http://pubs.acs.org.

## Author Contributions

The manuscript was written through contributions of all authors. All authors have given approval to the final version of the manuscript.

## Notes

M.R.A., L.B. and C.O. are founders of Ambagon Therapeutics. M.R.A is a director, L.B. is a member of Ambagon’s scientific advisory board, C.O., E.S are employees of Ambagon.

## ACKNOWLEDGMENTS

We thank the Renslo laboratory as well as Paul Burroughs and Joshua Ramos for the synthesis of the disulfide library; we also thank Julia Davies, Grace Pohan, and Amanda Paulson for the automated MS data processing infrastructure in the SMDC and compound data clustering. We thank Markella Konstantinidou and Priya Jaishankar for support with synthetic strategy and John Morrow for contributions in the initial stage of the 14-3-3 tethering project.

## Insert Table of Contents artwork here

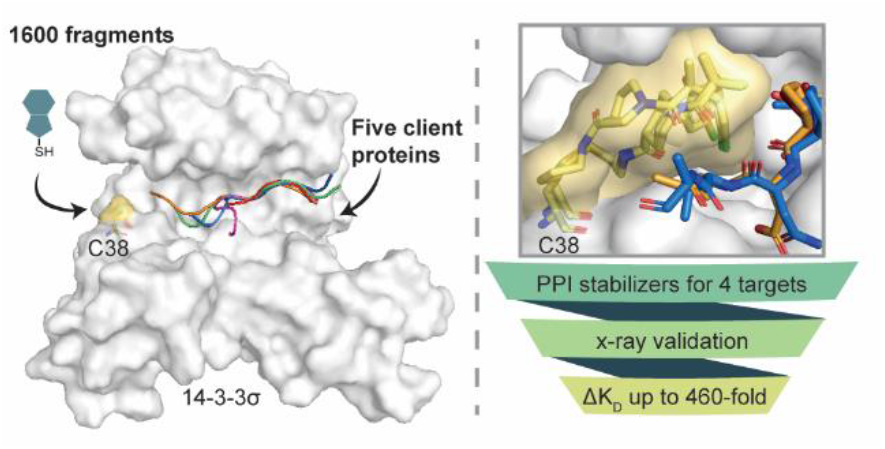

## References

(1) Garlick, J. M.; Mapp, A. K. Selective Modulation of Dynamic Protein Complexes. Cell Chem. Biol. 2020, 27 (8), 986–997. https://doi.org/10.1016/j.chembiol.2020.07.019.

(2) Ivanov, A. A.; Revennaugh, B.; Rusnak, L.; Gonzalez-Pecchi, V.; Mo, X.; Johns, M. A.; Du, Y.; Cooper, L. A. D.; Moreno, C. S.; Khuri, F. R.; Fu, H. The OncoPPi Portal: An Integrative Resource to Explore and Prioritize Protein-Protein Interactions for Cancer Target Discovery. Bioinforma. Oxf. Engl. 2018, 34 (7), 1183–1191. https://doi.org/10.1093/bioinformatics/btx743.

(3) Kim, M.; Park, J.; Bouhaddou, M.; Kim, K.; Rojc, A.; Modak, M.; Soucheray, M.; McGregor, M. J.; O’Leary, P.; Wolf, D.; Stevenson, E.; Foo, T. K.; Mitchell, D.; Herrington, K. A.; Muñoz, D. P.; Tutuncuoglu, B.; Chen, K. H.; Zheng, F.; Kreisberg, J. F.; Diolaiti, M. E.; Gordan, J. D.; Coppé, J. P.; Swaney, D. L.; Xia, B.; Veer, L. van ’t; Ashworth, A.; Ideker, T.; Krogan, N. J. A Protein Interaction Landscape of Breast Cancer. Science 2021, 374 (6563), abf3066. https://doi.org/10.1126/science.abf3066.

(4) Wright, M. T.; Plate, L. Revealing Functional Insights into ER Proteostasis through Proteomics and Interactomics. Exp. Cell Res. 2021, 399 (1), 112417. https://doi.org/10.1016/j.yexcr.2020.112417.

(5) Hu, G.; Wu, Z.; Uversky, V. N.; Kurgan, L. Functional Analysis of Human Hub Proteins and Their Interactors Involved in the Intrinsic Disorder-Enriched Interactions. Int. J. Mol. Sci. 2017, 18 (12), E2761. https://doi.org/10.3390/ijms18122761.

(6) Arkin, M. R.; Tang, Y.; Wells, J. A. Small-Molecule Inhibitors of Protein-Protein Interactions: Progressing toward the Reality. Chem. Biol. 2014, 21 (9), 1102–1114. https://doi.org/10.1016/j.chembiol.2014.09.001.

(7) Celis, S.; Hobor, F.; James, T.; Bartlett, G. J.; Ibarra, A. A.; Shoemark, D. K.; Hegedüs, Z.; Hetherington, K.; Woolfson, D. N.; Sessions, R. B.; Edwards, T. A.; Andrews, D. M.; Nelson, A.; Wilson, A. J. Query-Guided Protein-Protein Interaction Inhibitor Discovery. Chem. Sci. 2021, 12 (13), 4753–4762. https://doi.org/10.1039/d1sc00023c.

(8) Linhares, B. M.; Grembecka, J.; Cierpicki, T. Targeting Epigenetic Protein-Protein Interactions with Small-Molecule Inhibitors. Future Med. Chem. 2020, 12 (14), 1305–1326.

(9) Modell, A. E.; Blosser, S. L.; Arora, P. S. Systematic Targeting of Protein-Protein Interactions. Trends Pharmacol. Sci. 2016, 37 (8), 702–713. https://doi.org/10.1016/j.tips.2016.05.008.

(10) Zhong, M.; Lee, G. M.; Sijbesma, E.; Ottmann, C.; Arkin, M. R. Modulating Protein-Protein Interaction Networks in Protein Homeostasis. Curr. Opin. Chem. Biol. 2019, 50, 55–65. https://doi.org/10.1016/j.cbpa.2019.02.012.

(11) Andrei, S. A.; Sijbesma, E.; Hann, M.; Davis, J.; O’Mahony, G.; Perry, M. W. D.; Karawajczyk, A.; Eickhoff, J.; Brunsveld, L.; Doveston, R. G.; Milroy, L.-G.; Ottmann, C. Stabilization of Protein-Protein Interactions in Drug Discovery. Expert Opin. Drug Discov. 2017, 12 (9), 925–940. https://doi.org/10.1080/17460441.2017.1346608.

(12) Tang, C.; Mo, X.; Niu, Q.; Wahafu, A.; Yang, X.; Qui, M.; Ivanov, A. A.; Du, Y.; Fu, H. Hypomorph Mutation-Directed Small-Molecule Protein-Protein Interaction Inducers to Restore Mutant SMAD4-Suppressed TGF-β Signaling. Cell Chem. Biol. 2021, 28 (5), 636-647.e5. https://doi.org/10.1016/j.chembiol.2020.11.010.

(13) Stevers, L. M.; Sijbesma, E.; Botta, M.; MacKintosh, C.; Obsil, T.; Landrieu, I.; Cau, Y.; Wilson, A. J.; Karawajczyk, A.; Eickhoff, J.; Davis, J.; Hann, M.; O’Mahony, G.; Doveston, R. G.; Brunsveld, L.; Ottmann, C. Modulators of14-3-3 Protein–Protein Interactions. J. Med. Chem. 2018, 61 (9), 3755–3778. https://doi.org/10.1021/acs.jmed-chem.7b00574.

(14) De Vries-van Leeuwen, I. J.; da Costa Pereira, D.; Flach, K. D.; Piersma, S. R.; Haase, C.; Bier, D.; Yalcin, Z.; Michalides, R.; Feenstra, K. A.; Jiménez, C. R.; de Greef, T. F. A.; Brunsveld, L.; Ottmann, C.; Zwart, W.; de Boer, A. H. Interaction of 14-3-3 Proteins with the Estrogen Receptor Alpha F Domain Provides a Drug Target Interface. Proc. Natl. Acad. Sci. U. S. A. 2013, 110 (22), 8894–8899. https://doi.org/10.1073/pnas.1220809110.

(15) Mayor-Ruiz, C.; Bauer, S.; Brand, M.; Kozicka, Z.; Siklos, M.; Imrichova, H.; Kaltheuner, I. H.; Hahn, E.; Seiler, K.; Koren, A.; Petzold, G.; Fellner, M.; Bock, C.; Müller, A. C.; Zuber, J.; Geyer, M.; Thomä, N. H.; Kubicek, S.; Winter, G. E. Rational Discovery of Molecular Glue Degraders via Scalable Chemical Profiling. Nat. Chem. Biol. 2020, 16 (11), 1199–1207. https://doi.org/10.1038/s41589-020-0594-x.

(16) Chamberlain, P. P.; Cathers, B. E. Cereblon Modulators: Low Molecular Weight Inducers of Protein Degradation. Drug Discov. Today Technol. 2019, 31, 29–34. https://doi.org/10.1016/j.ddtec.2019.02.004.

(17) Słabicki, M.; Kozicka, Z.; Petzold, G.; Li, Y.-D.; Manojkumar, M.; Bunker, R. D.; Donovan, K. A.; Sievers, Q. L.; Koeppel, J.; Suchyta, D.; Sperling, A. S.; Fink, E. C.; Gasser, J. A.; Wang, L. R.; Corsello, S. M.; Sellar, R. S.; Jan, M.; Gillingham, D.; Scholl, C.; Fröhling, S.; Golub, T. R.; Fischer, E. S.; Thomä, N. H.; Ebert, B. L. The CDK Inhibitor CR8 Acts as a Molecular Glue Degrader That Depletes Cyclin K. Nature 2020, 585 (7824), 293–297. https://doi.org/10.1038/s41586-020-2374-x.

(18) Hartman, A. M.; Elgaher, W. A. M.; Hertrich, N.; Andrei, S. A.; Ottmann, C.; Hirsch, A. K. H. Discovery of Small-Molecule Stabilizers of 14-3-3 Protein-Protein Interactions via Dynamic Combinatorial Chemistry. ACS Med. Chem. Lett. 2020, 11 (5), 1041–1046. https://doi.org/10.1021/acsmedchemlett.9b00541.

(19) Pfaff, S. J.; Chimenti, M. S.; Kelly, M. J. S.; Arkin, M. R. Biophysical Methods for Identifying Fragment-Based Inhibitors of Protein-Protein Interactions. In Protein-Protein Interactions: Methods and Applications; Meyerkord, C. L., Fu, H., Eds.; Methods in Molecular Biology; Springer: New York, NY, 2015; pp 587–613. https://doi.org/10.1007/978-1-4939-2425-7_39.

(20) Erlanson, D. A.; Arndt, J. W.; Cancilla, M. T.; Cao, K.; Elling, R. A.; English, N.; Friedman, J.; Hansen, S. K.; Hession, C.; Joseph, I.; Kumaravel, G.; Lee, W.-C.; Lind, K. E.; McDowell, R. S.; Miatkowski, K.; Nguyen, C.; Nguyen, T. B.; Park, S.; Pathan, N.; Penny, D. M.; Romanowski, M. J.; Scott, D.; Silvian, L.; Simmons, R. L.; Tangonan, B. T.; Yang, W.; Sun, L. Discovery of a Potent and Highly Selective PDK1 Inhibitor via Fragment-Based Drug Discovery. Bioorg. Med. Chem. Lett. 2011, 21 (10), 3078–3083. https://doi.org/10.1016/j.bmcl.2011.03.032.

(21) Hallenbeck, K. K.; Davies, J. L.; Merron, C.; Ogden, P.; Sijbesma, E.; Ottmann, C.; Renslo, A. R.; Wilson, C.; Arkin, M. R. A Liquid Chromatography/Mass Spectrometry Method for Screening Disulfide Tethering Fragments. SLAS Discov. Adv. Life Sci. R D 2018, 23 (2), 183–192. https://doi.org/10.1177/2472555217732072.

(22) Raimundo, B. C.; Oslob, J. D.; Braisted, A. C.; Hyde, J.; McDowell, R. S.; Randal, M.; Waal, N. D.; Wilkinson, J.; Yu, C. H.; Arkin, M. R. Integrating Fragment Assembly and Biophysical Methods in the Chemical Advancement of Small-Molecule Antagonists of IL-2: An Approach for Inhibiting Protein-Protein Interactions. J. Med. Chem. 2004, 47 (12), 3111–3130. https://doi.org/10.1021/jm049967u.

(23) Lodge, J. M.; Rettenmaier, T. J.; Wells, J. A.; Pomerantz, W. C.; Mapp, A. K. FP Tethering: A Screening Technique to Rapidly Identify Compounds That Disrupt Protein-Protein Interactions. MedChemComm 2014, 5, 370–375. https://doi.org/10.1039/C3MD00356F.

(24) Sadowsky, J. D.; Burlingame, M. A.; Wolan, D. W.; McClendon, C. L.; Jacobson, M. P.; Wells, J. A. Turning a Protein Kinase on or off from a Single Allosteric Site via Disulfide Trapping. Proc. Natl. Acad. Sci. U. S. A. 2011, 108(15), 6056–6061. https://doi.org/10.1073/pnas.1102376108.

(25) Sijbesma, E.; Hallenbeck, K. K.; Leysen, S.; de Vink, P. J.; Skóra, L.; Jahnke, W.; Brunsveld, L.; Arkin, M. R.; Ottmann, C. Site-Directed Fragment-Based Screening for the Discovery of Protein–Protein Interaction Stabilizers. J. Am. Chem. Soc. 2019, 141 (8), 3524–3531. https://doi.org/10.1021/jacs.8b11658.

(26) Burlingame, M. A.; Tom, C. T. M. B.; Renslo, A. R. Simple One-Pot Synthesis of Disulfide Fragments for Use in Disulfide-Exchange Screening. ACS Comb. Sci. 2011, 13 (3), 205–208. https://doi.org/10.1021/co200038g.

(27) Morrison, D. K. The 14-3-3 Proteins: Integrators of Diverse Signaling Cues That Impact Cell Fate and Cancer Development. Trends Cell Biol. 2009, 19 (1), 16–23. https://doi.org/10.1016/j.tcb.2008.10.003.

(28) Obsilova, V.; Obsil, T. The 14-3-3 Proteins as Important Allosteric Regulators of Protein Kinases. Int. J. Mol. Sci. 2020, 21 (22), E8824. https://doi.org/10.3390/ijms21228824.

(29) Aghazadeh, Y.; Papadopoulos, V. The Role of the 14-3-3 Protein Family in Health, Disease, and Drug Development. Drug Discov. Today 2016, 21 (2), 278–287. https://doi.org/10.1016/j.drudis.2015.09.012

(30) Gardino, A. K.; Smerdon, S. J.; Yaffe, M. B. Structural Determinants of 14-3-3 Binding Specificities and Regulation of Subcellular Localization of 14-3-3-Ligand Complexes: A Comparison of the X-Ray Crystal Structures of All Human 14-3-3 Isoforms. Semin. Cancer Biol. 2006, 16 (3), 173–182. https://doi.org/10.1016/j.semcancer.2006.03.007.

(31) Zhao, J.; Meyerkord, C. L.; Du, Y.; Khuri, F. R.; Fu, H. 14-3-3 Proteins as Potential Therapeutic Targets. Semin. Cell Dev. Biol. 2011, 22 (7), 705–712. https://doi.org/10.1016/j.semcdb.2011.09.012.

(32) Molzan, M.; Kasper, S.; Röglin, L.; Skwarczynska, M.; Sassa, T.; Inoue, T.; Breitenbuecher, F.; Ohkanda, J.; Kato, N.; Schuler, M.; Ottmann, C. Stabilization of Physical RAF/14-3-3 Interaction by Cotylenin A as Treatment Strategy for RAS Mutant Cancers. ACS Chem. Biol. 2013, 8 (9), 1869–1875. https://doi.org/10.1021/cb4003464.

(33) Bank, R. P. D. RCSB PDB: Homepage. https://www.rcsb.org/ (accessed 2022-06-21).

(34) Liau, N. P. D.; Wendorff, T. J.; Quinn, J. G.; Steffek, M.; Phung, W.; Liu, P.; Tang, J.; Irudayanathan, F. J.; Izadi, S.; Shaw, A. S.; Malek, S.; Hymowitz, S. G.; Sudhamsu, J. Negative Regulation of RAF Kinase Activity by ATP Is Overcome by 14-3-3-Induced Dimerization. Nat. Struct. Mol. Biol. 2020, 27 (2), 134–141. https://doi.org/10.1038/s41594-019-0365-0.

(35) Ballone, A.; Lau, R. A.; Zweipfenning, F. P. A.; Ottmann, C. A New Soaking Procedure for X-Ray Crystallographic Structural Determination of Protein-Peptide Complexes. Acta Crystallogr. Sect. F Struct. Biol. Commun. 2020, 76 (Pt 10), 501–507. https://doi.org/10.1107/S2053230X2001122X.

(36) Saha, M.; Carriere, A.; Cheerathodi, M.; Zhang, X.; Lavoie, G.; Rush, J.; Roux, P. P.; Ballif, B. A. RSK Phosphorylates SOS1 Creating 14-3-3-Docking Sites and Negatively Regulating MAPK Activation. Biochem. J. 2012, 447 (1), 159–166. https://doi.org/10.1042/BJ20120938.

(37) Saline, M.; Badertscher, L.; Wolter, M.; Lau, R.; Gunnarsson, A.; Jacso, T.; Norris, T.; Ottmann, C.; Snijder, A. AMPK and AKT Protein Kinases Hierarchically Phosphorylate the N-Terminus of the FOXO1 Transcription Factor, Modulating Interactions with 14-3-3 Proteins. J. Biol. Chem. 2019, 294 (35), 13106–13116. https://doi.org/10.1074/jbc.RA119.008649.

(38) Centorrino, F.; Ballone, A.; Wolter, M.; Ottmann, C. Biophysical and Structural Insight into the USP8/14-3-3 Interaction. FEBS Lett. 2018, 592 (7), 1211–1220. https://doi.org/10.1002/1873-3468.13017.

(39) Molzan, M.; Schumacher, B.; Ottmann, C.; Baljuls, A.; Polzien, L.; Weyand, M.; Thiel, P.; Rose, R.; Rose, M.; Kuhenne, P.; Kaiser, M.; Rapp, U. R.; Kuhlmann, J.; Ottmann, C. Impaired Binding of 14-3-3 to C-RAF in Noonan Syndrome Suggests New Approaches in Diseases with Increased Ras Signaling. Mol. Cell. Biol. 2010, 30 (19), 4698–4711. https://doi.org/10.1128/MCB.01636-09.

(40) Falcicchio, M.; Ward, J. A.; Chothia, S. Y.; Basran, J.; Mohindra, A.; Macip, S.; Roversi, P.; Doveston, R. G. Cooperative Stabilisation of 14-3-3σ Protein–Protein Interactions via Covalent Protein Modification. Chem. Sci. 2021. https://doi.org/10.1039/D1SC02120F.

(41) Molzan, M.; Ottmann, C. Synergistic Binding of the Phosphorylated S233- and S259-Binding Sites of C-RAF to One 14-3-3ζ Dimer. J. Mol. Biol. 2012, 423 (4), 486–495. https://doi.org/10.1016/j.jmb.2012.08.009.

(42) Cossar, P. J.; Wolter, M.; van Dijck, L.; Valenti, D.; Levy, L. M.; Ottmann, C.; Brunsveld, L. Reversible Covalent Imine-Tethering for Selective Stabilization of 14-3-3 Hub Protein Interactions. J. Am. Chem. Soc. 2021, 143 (22), 8454–8464. https://doi.org/10.1021/jacs.1c03035.

