## Supplemental Information for "A Systematic Approach to the Discovery of Protein-Protein Interaction Stabilizers"

**Table of Contents**

Supplementary Methods……S2

- Protein expression and purification
- Peptide sequences
- Disulfide tethering and screen and data processing
- Dose response LC/MS experiments
- Dose response fluorescence anisotropy measurements
- X-ray crystallography

Supplementary Tables……S4

- Table S1 – S7

Supplementary Figures……S11

- Figure S1 – S19

Supporting References……S24

### Supplementary Methods

#### *Protein Expression and Purification*

The 14-3-3 σ isoform with a truncated C-terminus after T231 (ΔC; to enhance crystallization) and an N-terminal His6 tag was expressed in RosettaTM 2(DE3)pLysS competent E. coli (Novagen) from a pPROEX HTb expression vector. After transformation following manufacturer's instructions, single colonies were picked to inoculate 30 mL precultures (LB), which were added to 1.5 L terrific broth (TB) medium after overnight growth at 37 °C, 250 rpm. Expression was induced upon reaching OD600 1.9−2.1 by adding 400 μM IPTG. After overnight expression at 30°C, 150 rpm, cells were harvested by centrifugation at 6,500 rpm, resuspended in lysis buffer (50 mM HEPES pH 7.5, 500 mM NaCl, 20 mM imidazole, 10% glycerol, 1 mM TCEP), and lysed by sonication. The His6-tagged protein was first purified by Ni-affinity chromatography (Ni-NTA Agarose, Invitrogen) (Wash buffer 50 mM HEPES pH 7.5, 500 mM NaCl, 20 mM imidazole, 1 mM TCEP; Elution buffer 50 mM HEPES pH 7.5, 500 mM NaCl, 500 mM imidazole, 1 mM TCEP) followed by His-tag cleavage by TEV protease during dialysis (50 mM HEPES pH 7.5, 150 mM NaCl, 1 mM TCEP) overnight at 4 °C. The flowthrough of a second Ni-affinity column was collected and analyzed for purity by SDS-PAGE and Q-Tof LC/MS. The protein was buffer exchanged (Storage buffer 25 mM HEPES pH 7.5, 150 mM NaCl, 1 mM TCEP) and concentrated to ~13 mg/mL and aliquots flash-frozen for storage at −80°C. 14-3-3σΔC used for crystallography was concentrated to ~70 mg/mL and aliquots were flash-frozen for storage at -80 °C.

#### *Peptide Sequences*

Peptides for disulfide trapping were ordered from ScenicBio (Houston, Texas) as lyophilized solids. Sequences were as follows: ERα-pp (Ac-KYYITGEAEGF-PA{pT}V-COOH); SOS1-pp (Ac-PRRRPE{pS}APAESS-CONH2); USP8-pp (Ac-KLKRSY{pS}SPDITQ-CONH2); CRAF-pp (Ac-RQRST{pS}TPNVH-CONH2); FOXO1-pp (Ac-RPRSC{pT}WPLPR-CONH2. 2 mM peptide stocks were suspended in water. Fluorescein-labeled peptides were ordered from ScenicBio and Elim Biopharmaceuticals, Inc. (Hayward, CA) as lyophilized solids. Sequences were as written above with N-terminal 5-FAM-label for ERα-pp, SOS1-pp, USP8-pp, and FOXO1-pp. C-RAF-pp sequence was as follows: (5-FAM-QHRYSTPHAFTFNTSSPSSEGSLSQRQRST{pS}TPNVH-CONH2). 20 µM stocks were suspended in water. Peptides for crystallography were ordered from Genscript Biotech Corp. Sequences were as follows: ERα-pp (Ac-AEGFPA{pT}V-COOH); USP8-pp (Ac-KLKRSY{pS}SPDITQ-CONH2); CRAF-pp (QRST{pS}TPNVHH-CONH2); FOXO1-pp (Ac-RPR{pS}C{pT}WPLPR-CONH2). 5 mM stocks were suspended in Complexation Buffer (25 mM HEPES pH 7.5, 2 mM $MgCl_2$ and 2 mM βME).

#### *Disulfide Tethering Screen and Data Processing*

The primary disulfide tethering screen was performed by incubating target 14-3-3σ protein/phospho-peptide complex with small molecule in a 384-well plate format. The UCSF Small Molecule Discovery Center (SMDC) custom 1600 disulfide-containing fragment library was available as 50 mM stock solutions in DMSO. The screen was performed using 100 nM 14-3-3σ ΔC protein diluted in buffer (10 mM Tris, 250 μM betamercaptoethanol (βME), pH 8.0) and plated in 384-well plates (25 µL/well). 100 nL of each fragment was pinned from library master plates using non-sterile disposable 384 polypropylene pin tools (V & P Scientific) to give a final concentration of 200 µM. The peptide screens additionally contained a concentration of peptide equivalent to twice the $K_D$ as established by fluorescence anisotropy and competition experiments: (2 µM ERα-pp, 10 µM SOS1-pp, 10 µM USP8-pp, 18 µM CRAF-pp, and 750 nM FOXO1-pp). The reactions were incubated at room temperature for 3 hours before being measured by LC/MS (I-class Acquity UPLC/ Xevo G2-XS Quadrupole Time of Flight mass spectrometer, Waters). Data collection and automated processing followed a custom workflow, as previously described.[1] Compound resynthesis was done following the published procedures.[2]

#### *Dose Response LC/MS Experiments*

The initial mass spectrometry dose response follow-up experiments were performed under the same conditions as the primary screen with the exception that the compounds were titrated from 50 mM to 23 μM in a 3-fold dilution series in DMSO. Then 1 µL of the compound was transferred into 24 µL of protein-peptide solution for final concentrations of 2 mM – 914 nM and 4% DMSO. A final well of DMSO without compound was used as a control at the end of every dilution series. Compounds which displayed dose dependent activity in the presence of phospho-peptide were taken forward into more extensive dose responses in order to determine $DR_{50}$ values. Fragments were titrated from 50 mM to 0.129 nM in a 3-fold dilution series in DMSO. Then 1 µL of the compound was transferred into 24 µL of protein-peptide solution for final concentrations of 2 mM – 5.16 pM and 4% DMSO. A final well of DMSO without compound was used as a control at the end of every dilution series. Initial validation was done in single replicate. Hits were taken forward to fluorescence anisotropy experiments. $DR_{50}$ curves were graphed and calculated using log(agonist) vs. response – Variable slope (four parameters) from GraphPad Prism version 9.0 for Mac, GraphPad Software, San Diego, California USA, www.graphpad.com.

*Dose Response Fluorescence Anisotropy Experiments*

The fluorescence anisotropy compound dose responses were performed using 100 nM fluorescein-labeled peptides (with the exception of FOXO1-pp which was 10 nM) and 14-3-3σ protein diluted in buffer (10 mM HEPES pH 8.0, 150 mM NaCl, 0.01% TWEEN-20). 14-3-3σ protein concentration was equivalent to approximately half of the KD determined by fluorescence anisotropy and competition experiments (C-RAF: 1 µM 14-3-3σ; ERα: 250 nM 14-3-3σ; FOXO1: 25 nM 14-3-3σ; USP8: 2.5 µM 14-3-3σ). Fragments were titrated from 50 mM to 11.9 nM in a 2-fold dilution series in DMSO. 1 µL of compound was transferred in triplicates into 24 µL of protein-peptide solution for final concentrations of 2 mM – 0.477 nM and 4% DMSO. A final well of DMSO without compound was used as a control at the end of the dilution series. A row of 5-FAM-labeled peptide alone was used as a lower limit and a row of 5-FAM-labeled peptide with 14-3-3 excess of the KD was used as an upper limit when calculating the EC50 of the fragments. $EC_{50}$ measurements were graphed and calculated using log(agonist) vs. response – Variable slope (four parameters) from GraphPad Prism version 9.0 for Mac, GraphPad Software, San Diego, California USA, www.graphpad.com. 14-3-3σ protein titrations were performed using 2-fold dilution of 14-3-3σ in buffer from starting concentration of 250 µM to 59.6 pM for C-RAF and USP8 and 25 µM starting concentration to 5.96 pM for FOXO1 and ERα. Final well contained only 5-FAM-labeled peptide and compound as a control. 5-FAM-peptides were used at 100 nM except FOXO1 which was at 10 nM. Fragments were added at a single concentration of 1 mM in order to maximize 14-3-3σ engagement. Apparent $K_D$ values of 14-3-3σ/phospho-peptide in presence of compounds were compared to $K_D$ values of DMSO control. Fold stabilization ($\Delta K_D$) was determined as the quotient of the apparent $K_D$ of the DMSO control over the apparent $K_D$ with compound. $EC_{50}$ and apparent $K_D$ measurements were graphed and calculated using log(agonist) vs. response – Variable slope (four parameters) from GraphPad Prism version 9.0 for Mac, GraphPad Software, San Diego, California USA, www.graphpad.com.

*X-ray Crystallography*

The 14-3-3σΔC protein, acetylated ERα, C-RAF and FOXO1 peptides and fragments (stock solution of 50mM in DMSO) were dissolved in complexation buffer (25 mM HEPES pH 7.5, 2 mM $MgCl_2$ and 2 mM βME) and mixed in a 1:2:2 molar stoichiometry (protein:peptide:fragment) at a final protein concentration of 12 mg/mL. The complex was set up for sitting-drop crystallization after overnight incubation at 4 °C, in a custom crystallization liquor (0.095 M HEPES (pH 7.1, 7.3, 7.5, 7.7), 0.19 M $CaCl_2$, 24-29 % PEG 400 and 5% (v/v) glycerol). Crystals grew within 10 – 14 days at 4 °C. Crystals were fished and flash-cooled in liquid nitrogen. X-ray diffraction (XRD) data were either collected at the Deutsche Elektronen-Synchrotron (DESY) PETRA III beamline P11, Hamburg, Germany (all datasets except for 14-3-3σ/C-RafpS259/Compound 1) or at the Diamond Light Source (DLS) beamline Io3, Oxfordshire, United Kingdom (14-3-3σ/C-RafpS259/Compound 1).
Automatic processing was done by DESY or DLS using XDS9[3] for data indexing and integration Initial processed data was then scaled using AIMLESS[4,5] and Molrep[6] was used for limited molecular replacement using PDB ID's 4JC3 and 3IQU as template. Presence of co-crystallized ligands was verified by visual inspection of the Fo-Fc and 2Fo-Fc electron density maps in Coot.[7] If electron density corresponding to the co-crystallized ligand was present, its structure and restrains were generated using eLBOW[8] before final model rebuilding and refinement was done using phenix.refine[9,10] and Coot. See Table S6 and S7 for data collection and refinement statistics. The structures were submitted to the PDB with IDs 8AFN, 8AV0, 8ADM, 8A62, 8A65, 8A68, 8A6H, 8A6F.

Supplementary Tables

**Table S1. Tethering and stabilization of 14-3-3/C-RAF primary screen hit compounds**

| | Primary Screen (250 µM BME) | MSDR (250 µM BME) | FADR (50 µM BME) | Protein Titrations (50 µM BME) | | |
|---|---|---|---|---|---|---|
| **SMDC ID** | **% Tethering** | **$DR_{50}$ (µM)** | **$EC_{50}$ (µM)** | **$K_{D\_app}$** | **$K_{D\_DMSO}$** | **Fold Stabilization** |
| 916851 (**7**) | 42.4 | 12.2 | 3.18 | 294 nM | 23 µM | 77 |
| 916856 (**8**) | 58.3 | 4.96 | 13.5 | 207 nM | 23 µM | 110 |
| 916935 | 44.9 | >2000 | — | — | — | — |
| 917041 | 38 | >2000 | — | — | — | — |
| 917072 | 62.8 | 465 | >2000 | — | — | — |
| 917215 | 40.4 | 5.01 | >2000 | — | — | — |
| 917215 | 38.3 | 5.01 | >2000 | — | — | — |
| 917285 | 56.7 | 5.61 | — | — | — | — |
| 917285 | 38.4 | 5.61 | — | — | — | — |
| 917323 | 42.5 | >2000 | — | — | — | — |
| 917426 | 40.9 | 5.76 | — | — | — | — |
| 917692 | 62.2 | >2000 | >2000 | — | — | — |
| 917725 | 42.5 | — | — | — | — | — |
| 917729 | 38.5 | 5.24 | — | — | — | — |
| 917746 | 35.4 | <0.91 | 19.4 | — | — | — |
| 917748 | 15.8+/-17 | >2000 | — | — | — | — |
| 917755 | 51.1 | 5.82 | >2000 | — | — | — |
| 917845 | 13.9+/-16 | — | — | — | — | — |
| 917866 | 52.2 | >2000 | >2000 | — | — | — |
| 917908 | 64.7 | >2000 | — | — | — | — |
| 917922 | 47.1 | >2000 | >2000 | — | — | — |
| 917949 (**1**) | 85.3 | 0.007 | 0.922 | 106 nM | 8.5 µM | 81 |
| 917963 | 52 | 5.29 | >2000 | — | — | — |
| 917982 | 63.7 | 5.45 | 0.58 | — | — | — |
| 917988 | 55.4 | >2000 | — | — | — | — |
| 917995 | 41.7 | >2000 | — | — | — | — |
| 917999 (**5**) | 37.1 | 0.58 | 0.22 | 92 | 23 µM | 246 |
| 918000 | 60.4 | 3.95 | — | — | — | — |
| 918039 (**6**) | 48.3 | 8.78 | 1.33 | 100 nM | 42 µM | 426 |

**Table S2. Tethering and stabilization of 14-3-3/FOXO1 primary screen hit compounds**

| | Primary Screen (250 μM BME) | MSDR (250 μM BME) | FADR (50 μM BME) | Protein Titrations (50 μM BME) | | |
|---|---|---|---|---|---|---|
| **SMDC ID** | **% Tethering** | **$DR_{50}$ (μM)** | **$EC_{50}$ (μM)** | **$K_{D\_app}$** | **$K_{D\_DMSO}$** | **Fold Stabilization** |
| 917034 | 29.7 | >2000 | — | — | — | — |
| 917041 | 34.3 | >2000 | — | — | — | — |
| 917222 | 50.6 | — | — | — | — | — |
| 917234 | 34.3 | 16.9 | — | — | — | — |
| 917261 | 57.5 | 3.61 | — | — | — | — |
| 917282 | 48.4 | — | — | — | — | — |
| 917316 | 83.6 | 0.042 | — | 117 nM | 87 nM | — |
| 917346 | 36.5 | >2000 | — | — | — | — |
| 917383 (**4**) | 34.3 | 143.4 | 6.69 | 9.7 nM | 111 nM | 12 |
| 917429 | 28.5 | >2000 | — | — | — | — |
| 917607 | 35.4 | — | — | — | — | — |
| 917701 (**3**) | 68.7 | 2.2 | 0.85 | 10.6 nM | 42 nM | 4 |
| 917720 | 31.4 | >2000 | — | — | — | — |
| 917726 (**2**) | 67.6 | 0.36 | 5.1 | 7.7 nM | 39 nM | 5 |
| 917748 | 14.8+/-14 | >2000 | — | — | — | — |
| 917773 | 47.5 | >2000 | — | — | — | — |
| 917800 | 66.6 | 4.78 | 2.41 | 69 nM | 66 nM | — |
| 917845 | 13.8+/-14 | — | — | — | — | — |
| 917848 | 10.4+/-13 | >2000 | — | — | — | — |
| 917922 | 57.9 | >2000 | — | — | — | — |
| 917967 | 31.2 | >2000 | — | — | — | — |
| 917986 | 38 | >2000 | — | — | — | — |
| 917992 | 38.9 | 48.7 | — | — | — | — |
| 957764 | 30.7 | >2000 | — | — | — | — |
| 957769 | 38.6 | >2000 | — | — | — | — |
| 960037 | 31.9 | — | — | — | — | — |
| 966848 | 28 | >2000 | — | — | — | — |
| 966851 | 37.1 | >2000 | — | — | — | — |
| 966864 | 43.4 | >2000 | — | — | — | — |
| 966943 | 40.2 | >2000 | — | — | — | — |

**Table S3. Tethering and stabilization of 14-3-3/ERα primary screen hit compounds**

| | Primary Screen (250 μM BME) | MSDR (250 μM BME) | FADR (50 μM BME) | Protien Titrations (50 μM BME) | | |
|---|---|---|---|---|---|---|
| **SMDC ID** | **% Tethering** | **$DR_{50}$ (μM)** | **$EC_{50}$ (μM)** | **$K_{D\_app}$** | **$K_{D\_DMSO}$** | **Fold Stabilization** |
| 916848 | 40.9 | >2000 | >2000 | — | — | — |
| 916854 | 35.1 | 786 | — | — | — | — |
| 916856 (**8**) | 35.7 | 14.5 | — | — | — | — |
| 916874 | 47.3 | >2000 | — | — | — | — |
| 916933 | 45.6 | >2000 | — | — | — | — |
| 916935 | 43 | >2000 | — | — | — | — |
| 916936 | 33.3 | >2000 | — | — | — | — |
| 916945 | 56.9 | >2000 | — | — | — | — |
| 916974 | 34.9 | 133 | — | — | — | — |
| 917101 | 38.1 | 74.1 | — | — | — | — |
| 917607 | 40.5 | 1900 | >2000 | — | — | — |
| 917692 | 47.3 | >2000 | — | — | — | — |
| 917717 | 36.7 | >2000 | >2000 | — | — | — |
| 917725 | 51.5 | >2000 | — | — | — | — |
| 917730 | 37.6 | >2000 | 1.51 | — | — | — |
| 917748 | 20.8+/-17 | 18.5 | 129 | — | — | — |
| 917750 | 18.3+/-14 | >2000 | >2000 | — | — | — |
| 917770 | 17.3+/-13 | >2000 | — | — | — | — |
| 917848 | 17.2+/-13 | >2000 | >2000 | — | — | — |
| 917878 | 36.1 | >2000 | — | — | — | — |
| 917922 | 73.1 | 3.67 | — | — | — | — |
| 917949 (**1**) | 73.1 | 18.1 | 1.31 | 21 nM | 360 nM | 19 |
| 917952 | 33.4 | >2000 | — | — | — | — |
| 917982 | 37.1 | >2000 | — | — | — | — |
| 918016 | 35.7 | 4.47 | — | — | — | — |
| 957766 | 44.9 | 459 | — | — | — | — |
| 957779 | 63.6 | >2000 | — | — | — | — |
| 960009 | 49.4 | — | — | — | — | — |
| 960037 | 56.2 | 8.31 | — | — | — | — |
| 966784 | 33.8 | >2000 | — | — | — | — |
| 966840 | 43 | >2000 | — | — | — | — |
| 993941 | 34.2 | — | — | — | — | — |
| 994336 | 38.2 | — | — | — | — | — |
| 994825 | 39.8 | — | — | — | — | — |

**Table S4. Tethering and stabilization of 14-3-3/USP8 primary screen hit compounds**

| | Primary Screen (250 μM BME) | MSDR (250 μM BME) | FADR (50 μM BME) | Protein Titrations (50 μM BME) | | |
|---|---|---|---|---|---|---|
| **SMDC ID** | **% Tethering** | **$DR_{50}$ (μM)** | **$EC_{50}$ (μM)** | **$K_{D\_app}$** | **$K_{D\_DMSO}$** | **Fold Stabilization** |
| 916848 | 0 | >2000 | 4.21 | — | — | — |
| 916933 | 38.3 | >2000 | — | — | — | — |
| 916945 | 36.5 | >2000 | — | — | — | — |
| 916974 | 39.4 | >2000 | — | — | — | — |
| 917041 | 30.1 | >2000 | >2000 | — | — | — |
| 917252 | 44.1 | >2000 | | — | — | — |
| 917340 | 50.4 | 648 | 1140 | — | — | — |
| 917644 | 18.7+/-23 | — | | — | — | — |
| 917659 | 28.9 | >2000 | >2000 | — | — | — |
| 917692 | 51.1 | 240 | — | — | — | — |
| 917748 | 19.5+/-19 | — | — | — | — | — |
| 917848 | 15.5+/-13 | >2000 | — | — | — | — |
| 917949 (**1**) | 52.3 | 0.024 | 3.38 | 1.1 | 4.5 | 4 |
| 957766 | 39.7 | 1636 | >2000 | — | — | — |
| 957779 | 59.8 | >2000 | >2000 | — | — | — |
| 957817 | 35.1 | >2000 | — | — | — | — |
| 957821 | 57.5 | — | — | — | — | — |
| 957824 | 36 | >2000 | — | — | — | — |
| 957999 | 29.4 | >2000 | — | — | — | — |
| 958001 | 53.9 | >2000 | >2000 | — | — | — |
| 960009 | 67.6 | — | — | — | — | — |
| 960031 | 12.8+/-13 | >2000 | — | — | — | — |
| 960037 | 52.9 | 1749 | >2000 | — | — | — |
| 967287 | 35.8 | — | — | — | — | — |
| 967289 | 30.4 | — | — | — | — | — |
| 994439 | 36.1 | — | — | — | — | — |
| 994487 | 41.1 | — | — | — | — | — |
| 994782 | 31 | — | — | — | — | — |
| 994821 | 48.2 | — | — | — | — | — |

**Table S5. Tethering and stabilization of 14-3-3/SOS1 primary screen hit compounds**

| | Primary Screen (250 µM BME) | MSDR (250 µM BME) | FADR (50 µM BME) | Prot Titrations (50 µM BME) | | |
|---|---|---|---|---|---|---|
| **SMDC ID** | **% Tethering** | **$DR_{50}$ (µM)** | **$EC_{50}$ (µM)** | **$K_{D\_app}$** | **$K_{D\_DMSO}$** | **Fold Stabilization** |
| 916848 | 28.3 | >2000 | 1598 | — | — | — |
| 916933 | 32.5 | >2000 | 297 | — | — | — |
| 916935 | 22.3 | — | — | — | — | — |
| 916992 | 28.4 | >2000 | >2000 | — | — | — |
| 917041 | 23 | >2000 | — | — | — | — |
| 917101 | 26.9 | >2000 | — | — | — | — |
| 917340 | 27.2 | >2000 | 55.1 | — | — | — |
| 917447 | 37.1 | >2000 | 148 | — | — | — |
| 917602 | 9.33+/-12 | >2000 | — | — | — | — |
| 917609 | 11.3+/-14 | >2000 | — | — | — | — |
| 917611 | 9.73+/-12 | >2000 | — | — | — | — |
| 917621 | 10.7+/-13 | >2000 | — | — | — | — |
| 917692 | 27.3 | 325 | >2000 | — | — | — |
| 917748 | 15.1+/-16 | 4.26 | — | — | — | — |
| 917845 | 9.67+/-12 | 266 | — | — | — | — |
| 917848 | 14.1+/-15 | >2000 | — | — | — | — |
| 917879 | 10.4+/-12 | >2000 | — | — | — | — |
| 917922 | 72.9 | 1912 | — | — | — | — |
| 917941 | 23.2 | >2000 | >2000 | — | — | — |
| 957779 | 54.1 | >2000 | — | — | — | — |
| 957795 | 55.3 | >2000 | — | — | — | — |
| 957999 | 37.1 | >2000 | — | — | — | — |
| 958001 | 67.8 | >2000 | — | — | — | — |
| 960009 | 50 | 28.2 | — | — | — | — |
| 960031 | 15.2+/-15 | >2000 | — | — | — | — |
| 960037 | 61.9 | 562 | 241 | — | — | — |
| 960039 | 7.73+/-10 | >2000 | — | — | — | — |
| 966837 | 26.3 | >2000 | — | — | — | — |
| 966840 | 55.9 | >2000 | — | — | — | — |

**Table S6. XRD data collection and refinement statistics for 14-3-3σ – Compound 1 structures**

| 14-3-3σ ΔC / Compound 1 | ERα | C-RAF | USP8 |
|---|---|---|---|
| **PDB ID** | **8AFN** | **8AV0** | **8ADM** |
| **Data collection** | | | |
| **Collection source** | DESY Petra III | Diamond Light Source I03 | DESY Petra III |
| **Collection date** | 2019-05-14 | 2019-08-08 | 2019-03-01 |
| **Wavelength (Å)** | 1.033 | 0.976 | 1.03320 |
| **Resolution (Å)** | 66.01 – 1.36 (1.38 – 1.36) | 34.31 – 1.50 (1.53 – 1.50) | 39.84 – 1.70 (1.73 – 1.70) |
| **Space group** | C2221 | C2221 | C2221 |
| **Unit cell** | 81.74 111.90 62.56 | 81.88 112.29 62.865 | 81.09 96.06 79.68 |
| **Total reflections**[a] | 113349 (4002) | 888809 (4361) | 452637 (22681) |
| **Unique reflections**[a] | 59489 (2234) | 46192 (2239) | 34318 (1669) |
| **Redundancy**[a] | 1.9 (1.8) | 1.9 (1.9) | 13.2 (13.6) |
| **Completeness (%)**[a] | 96.3 (69.3) | 99.1 (98.1) | 99.4 (98.5) |
| **Average I/σ(I)** [a] | 26.6 (1.7) | 53.1 (23.2) | 12.1 (2.5) |
| **Wilson B-factor (Å$^2$)** | 15.19 | 13.68 | 26.22 |
| **$CC_{1/2}$**[a,b,c] | 1.0 (0.68) | 1.0 (0.998) | 0.998 (0.832) |
| **$R_{merge}$**[a,c,d] | 0.015 (0.412) | 0.007 (0.024) | 0.113 (1.032) |
| **$R_{meas}$**[a,c,e] | 0.022 (0.582) | 0.011 (0.034) | 0.117 (1.072) |
| **Refinement** | | | |
| **Reflections in set: Refinement / R-free** | 59485 (4416) / 2967 (118) | 46186 (4524) / 2283 (227) | 34296 (3350) / 1762 (151) |
| **Non-H atoms: Overall / solvent** | 1882 / 320 | 2189 (240) | 1906 / 238 |
| **$R_{work}$ / $R_{free}$** | 0.1726 (0.2868) / 0.1844 (0.2899) | 0.1463 (0.0999) / 0.1802 (0.1582) | 0.1845 (0.2460) / 0.2197 (0.2903) |
| **RMSD from ideal geometry: Bond length (Å) / angles (°)** | 0.008 / 1.04 | 0.011 / 1.13 | 0.010 / 1.23 |
| **Average protein B-factor (Å$^2$)** | 20.71 | 20.53 | 33.18 |
| **Ramachandran: Favored / outlier (%)** | 98.30 / 0.00 | 98.73 (0.00) | 98.20 / 0.00 |
| **Clashscore** | 3.74 | 4.43 | 2.69 |

[a] Number in parentheses is for the highest resolution shell used in the refinement

[b] $CC_{1/2}$ = Pearson's intra-dataset correlation coefficient, as described by Karplus and Diederichs.[11]

[c] $R_{merge}$ (= $R_{sym}$) = $\sum_h \sum_1 |I_{h1} - \langle I_h \rangle| / \sum_h \sum_1 \langle I_h \rangle$, where $I_{h1}$ is the intensity of the 1th observation of reflection h and $\langle I_h \rangle$ is the average intensity of reflection h

[d] $R_{meas} = \sum_h \left| \sqrt{(n_h/(n_h-1)} \sum_1 |I_{h1} - \langle I_h \rangle| \right| / \sum_h \sum_1 \langle I_h \rangle$ where $n_h$ is the number of observations of reflection $h$

[e] Correlation of experimental intensities with intensities calculated from refined model, as described by Karplus and Diederichs.[11]

**Table S7. XRD data collection and refinement statistics for 14-3-3σ – FOXO1 and C-RAF structures**

| 14-3-3σ ΔC | Compound 2 / FOXO1 | Compound 3 / FOXO1 | Compound 5 / C-RAF | Compound 7 / C-RAF | Compound 8 / CRAF |
|---|---|---|---|---|---|
| **PDB ID** | **8A62** | **8A65** | **8A68** | **8A6H** | **8A6F** |
| **Data collection** | | | | | |
| **Collection source** | DESY Petra III | DESY Petra III | DESY Petra III | DESY Petra III | DESY Petra III |
| **Collection date** | 2021-11-04 | 2021-11-04 | 2022-03-24 | 2021-11-04 | 2021-11-04 |
| **Wavelength (Å)** | 1.033220 | 1.033220 | 1.033200 | 1.03320 | 1.03320 |
| **Resolution (Å)** | 45.41 – 1.60 (1.63 – 1.60) | 45.47 – 1.60 (1.63 – 1.60) | 45.62 – 1.60 (1.63 – 1.60) | 45.63 – 1.60 (1.63 – 1.60) | 45.77 – 1.60 (1.63 – 1.60) |
| **Space group** | C2221 | C2221 | C2221 | C2221 | C2221 |
| **Unit cell** | 81.60 111.50 62.71 | 81.88 112.14 62.63 | 82.40 112.26 62.75 | 82.50 112.23 62.76 | 82.94 112.91 62.79 |
| **Total reflections**[a] | 512777 (22392) | 518652 (22475) | 522241 (25556) | 522752 (22913) | 507054 (19971) |
| **Unique reflections**[a] | 37086 (1773) | 38393 (1881) | 38586 (1901) | 38619 (1878) | 39243 (1906) |
| **Redundancy**[a] | 13.8 (12.6) | 13.5 (11.9) | 13.5 (13.4) | 13.5 (12.2) | 12.9 (10.5) |
| **Completeness (%)**[a] | 97.6 (95.2) | 100.0 (99.9) | 99.6 (99.8) | 99.6 (98.8) | 100 (99.9) |
| **Average I/σ(I)** [a] | 39.0 (14.6) | 55.7 (19.9) | 38.8 (14.4) | 42.6 (14.7) | 14.6 (5.5) |
| **Wilson B-factor (Å²)** | 12.21 | 14.17 | 13.06 | 13.25 | 16.95 |
| **$CC_{1/2}$**[a,b,c] | 1.000 (0.992) | 1.000 (0.997) | 0.999 (0.995) | 1.0 (0.995) | 0.996 (0.437) |
| **$R_{merge}$**[a,c,e] | 0.044 (0.144) | 0.030 (0.102) | 0.044 (0.141) | 0.040 (0.115) | 0.221 (3.285) |
| **$R_{meas}$**[a,d,e] | 0.046 (0.150) | 0.031 (0.106) | 0.045 (0.147) | 0.041 (0.124) | 0.230 (3.472) |
| **Refinement** | | | | | |
| **Reflections in set: Refinement / R-free** | 37081 (3497) / 1895 (191) | 38370 (3794) / 1954 (201) | 38566 (3841) / 1962 (209) | 38596 (3798)/ 1965 (206) | 39110 (3778) / 1992 (192) |
| **Non-H atoms: Overall / solvent** | 2326 / 364 | 1918 / 342 | 1960 / 335 | 1934 / 355 | 1939 / 332 |
| **$R_{work}$ / $R_{free}$ (%)** | 0.1569 (0.1605) / 0.1819 (0.1933) | 0.1585 (0.1581) / 0.1807 (0.1929) | 0.1594 (0.1545) / 0.1791 (0.1725) | 0.1526 (0.1479) / 0.1737 (0.1853) | 0.1622 (0.1894) / 0.1810 (0.2209) |
| **RMSD from ideal geometry: Bond length (Å) / angles (°)** | 0.009 / 1.06 | 0.010 / 1.07 | 0.010 / 1.04 | 0.010 / 1.01 | 0.010 / 1.01 |
| **Average protein B-factor (Å²)** | 16.85 | 19.60 | 17.65 | 18.50 | 22.58 |
| **Ramachandran: Favored / outlier (%)** | 98.31 / 0.00 | 97.46 / 0.85 | 97.48 / 0.00 | 97.07 / 0.42 | 97.91 / 0.00 |
| **Clashscore** | 1.81 | 5.24 | 3.88 | 3.37 | 3.36 |

[a] Number in parentheses is for the highest resolution shell used in the refinement

[b] $CC_{1/2}$ = Pearson's intra-dataset correlation coefficient, as described by Karplus and Diederichs.[11]

[c] $R_{merge}$ (= $R_{sym}$) = $\sum_h \sum_1 |I_{h1} - < I_h >| / \sum_h \sum_1 < I_h >$, where $I_{h1}$ is the intensity of the 1th observation of reflection h and $< I_h >$ is the average intensity of reflection h

[d] $R_{meas}$ = $\sum_h \left| \sqrt{(n_h/(n_h - 1)} \sum_1 |I_{h1} - < I_h >| \right| / \sum_h \sum_1 < I_h >$ where $n_h$ is the number of observations of reflection $h$

[e] Correlation of experimental intensities with intensities calculated from refined model, as described by Karplus and Diederichs.[11]

Supplementary Figures

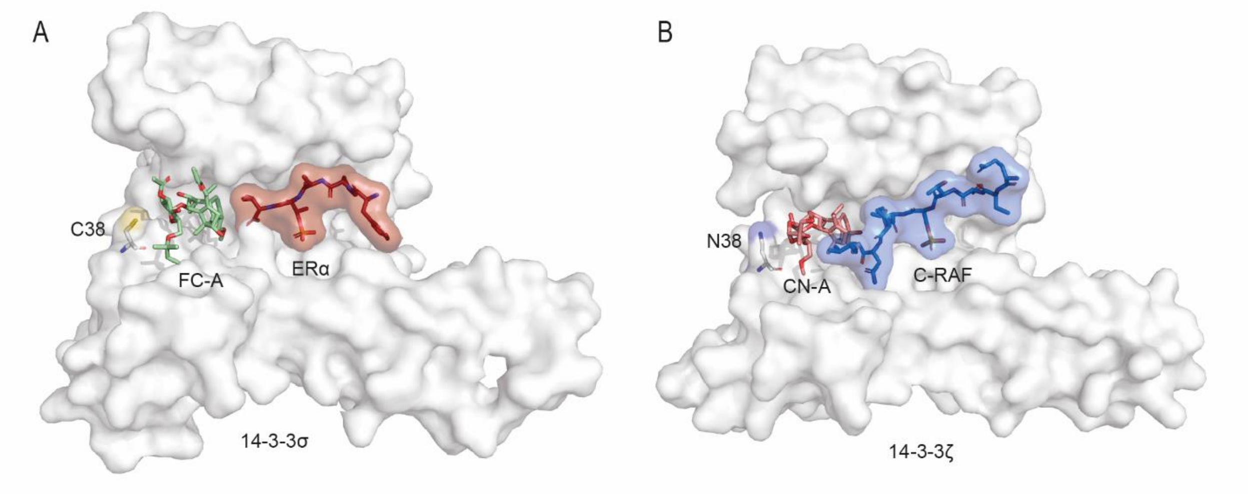


**Figure S1.** Crystal structures of natural product stabilizers (A) Fusicoccin-A (green sticks) and (B) Cotylenin-A (pinks sticks) bound to 14-3-3 (white surface) and clients ERα (red sticks) and C-RAF (blue sticks), respectively. Cysteine 38 depicted as white sticks. PDB ID: J4DD and 4IHL

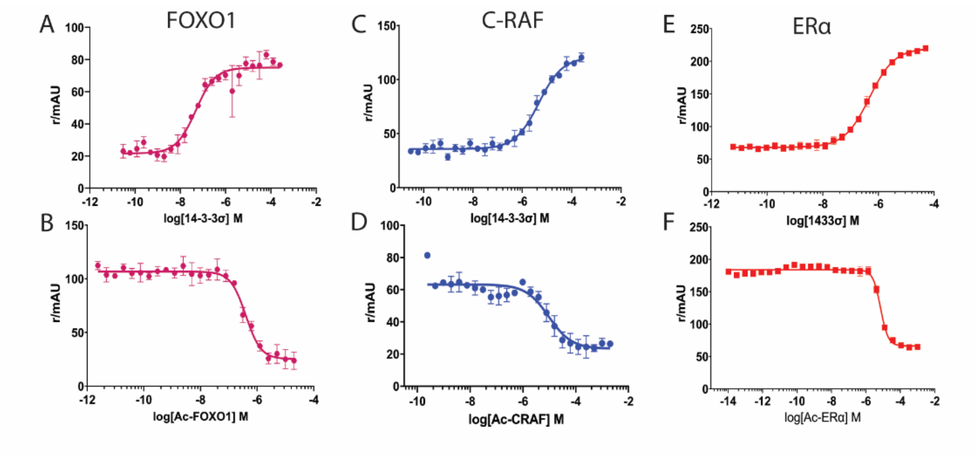


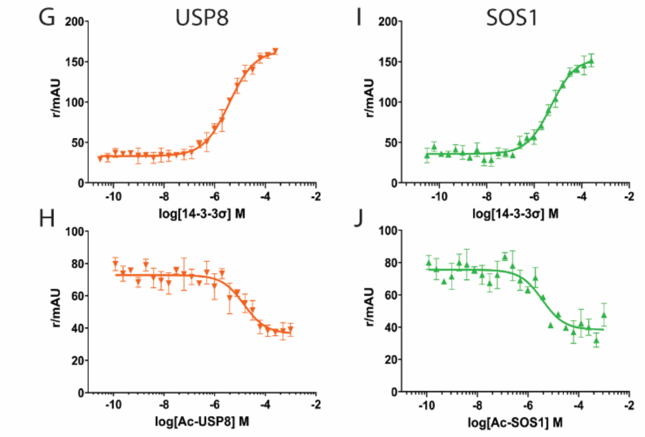


K

| Peptide | $K_D$ (14-3-3σ Titration) | $K_i$ (Ac-peptide competition) |
|---|---|---|
| FOXO1 | 48 nM | 396 nM |
| C-RAF | 4.64 µM | 9.50 µM |
| Erα | 476 nM | 7.10 µM |
| USP8 | 3.77 µM | 14.3 µM |
| SOS1 | 5.06 µM | 3.66 µM |

**Figure S2.** 14-3-3σ/phosphopeptide binding curves to determine binding affinities. (A) Protein titration with 14-3-3σ in presence of 100 nM FITC-labeled FOXO1 11-mer peptide determined $K_D$. (B) Competition assay used to determine $K_i$ of acetylated FOXO1 11-mer peptide (Ac-FOXO1) for mass spectrometry experiments. Ac-FOXO1 was titrated from high to low concentration in presence of 100 nM FITC-FOXO1 peptide and 14-3-3σ at 3/2 $K_D$ value determined in (A). (C) Protein titration with 14-3-3σ in presence of 100 nM FITC-labeled C-RAF 36-mer peptide determined $K_D$. (D) Competition assay used to determine $K_i$ of acetylated C-RAF 11-mer peptide (Ac-CRAF). (E) Protein titration with 14-3-3σ in presence of 100 nM FITC-labeled ERα 15-mer peptide determined $K_D$. (F) Competition assay used to determine $K_i$ of acetylated ERα 15-mer peptide (Ac-ERα). (G) Protein titration with 14-3-3σ in presence of 100 nM FITC-labeled USP8 13-mer peptide determined $K_D$. (H) Competition assay used to determine $K_i$ of acetylated USP8 13-mer peptide (Ac-USP8). (I) Protein titration with 14-3-3σ in presence of 100 nM FITC-labeled SOS1 13-mer peptide determined $K_D$. (J) Competition assay used to determine $K_i$ of acetylated SOS1 13-mer peptide (Ac-SOS1). (K) Table of peptide affinities to 14-3-3σ based on data from A-J.

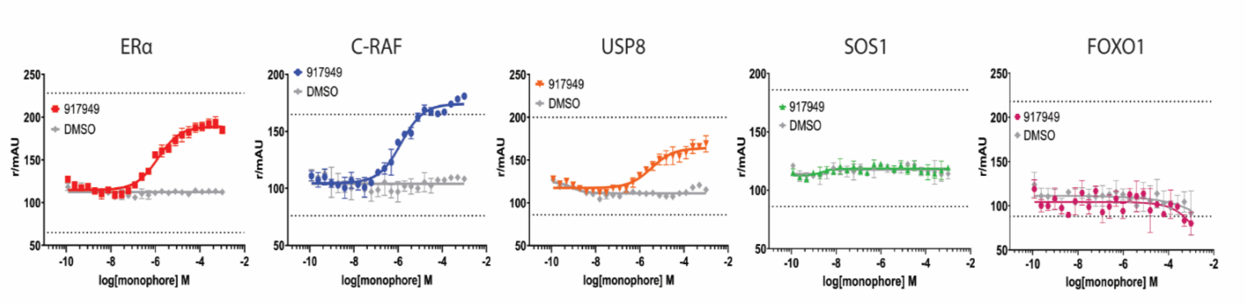


**Figure S3.** Fluorescence anisotropy dose response (FADR) curves for non-selective stabilizer 917949 (compound **1** in main text). Top black dashed line signifies maximum anisotropy control, the value of which was determined by peptide bound to 250 μM 14-3-3σ (at least 50-fold excess of peptide $K_D$). Bottom dashed line signifies minimum anisotropy control, value determined by peptide only and no 14-3-3σ. From left to right: 917949 dose-dependent stabilization of ERα phospho-peptide (red) binding to 14-3-3σ; 917949 stabilization of C-RAF (blue) binding to 14-3-3σ; 917949 stabilization of USP8 (orange) binding to 14-3-3σ. SOS1 (green) and FOXO1 (pink) showed no stabilization as compared to DMSO control (grey). From left to right $EC_{50}$ values: 1.31 μM (ERα); 922 nM (C-RAF); 3.38 μM (USP8); >2000 μM (SOS1); >2000 μM (FOXO1)

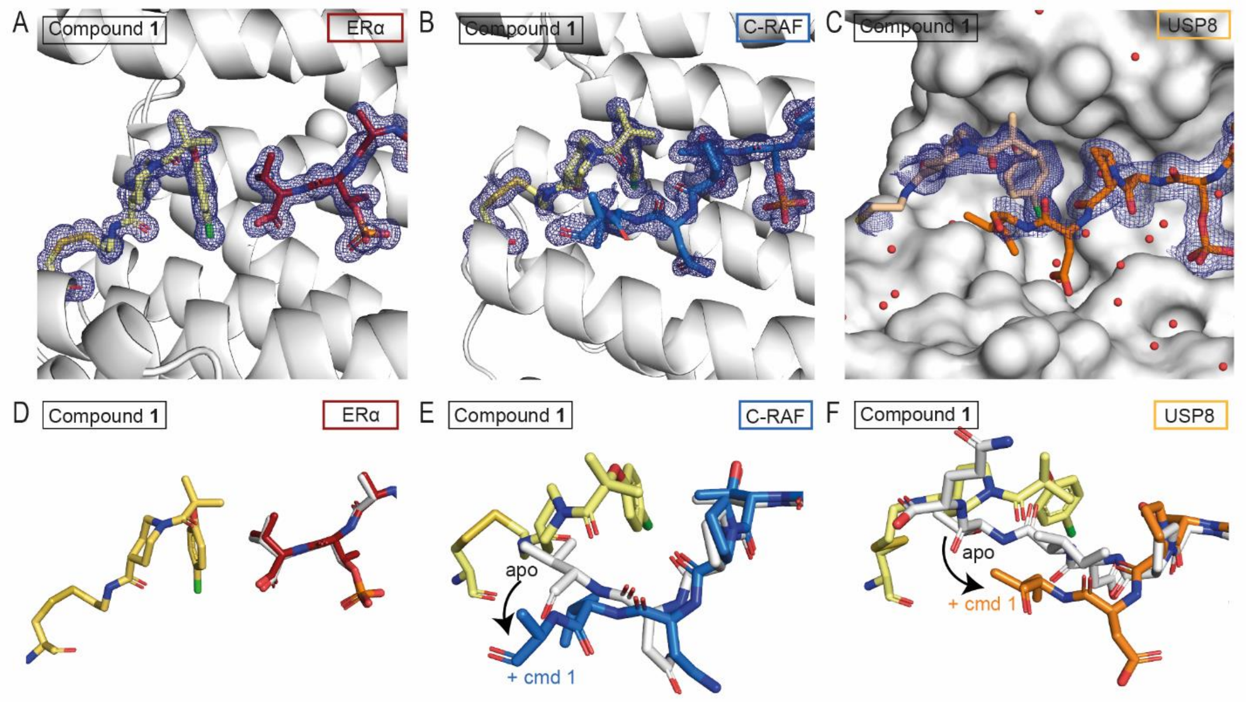


**Figure S4:** (A-C) Crystal structures of Compound **1** (yellow sticks) with (A) ERα (red sticks) (B) C-RAF (blue sticks) and USP8 (orange sticks) complexed with 14-3-3σ (white cartoon and surface). (D-F) Overlay of crystal structures of compound **1** (yellow sticks) – peptide complex with (D) ERα (red sticks when complexed with compound **1**, white sticks as apo), (E) C-RAF (blue sticks when complex with compound **1**, white sticks as apo), (F) USP8 (orange sticks when complexed with compound **1**, white sticks as apo). PDB ID apo structures: 4JC3, 4FJ3, 6F09. 2Fo-Fc electron density maps are contoured at 1σ.

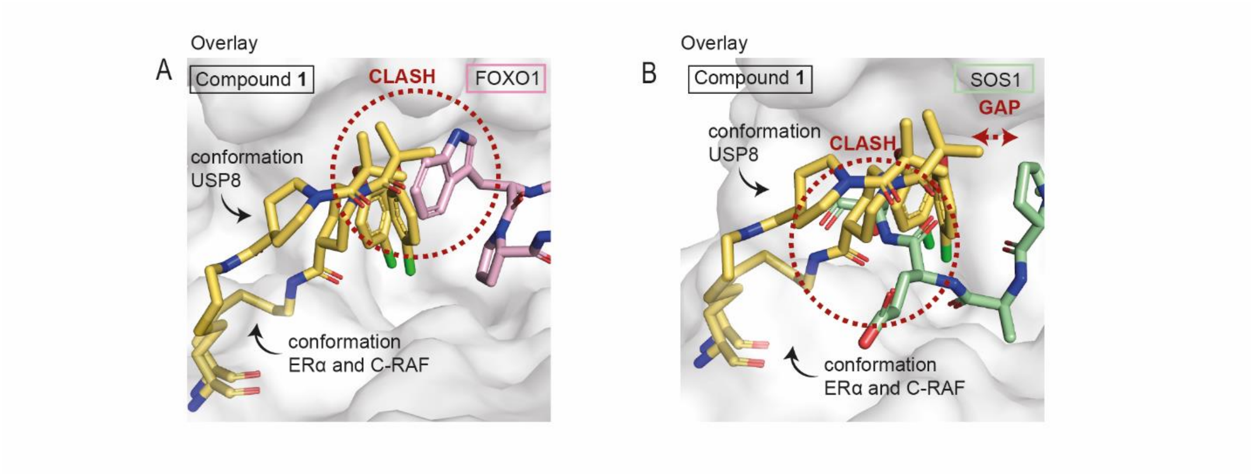


**Figure S5:** Crystallographic overlay of compound **1** (yellow sticks) with (A) FOXO1 (pink sticks) and (B) SOS1 (green sticks) in 14-3-3 (white surface), highlighting the clash of compound **1** with these peptides. PDB ID apo structures: 6QZR and 6Y44.

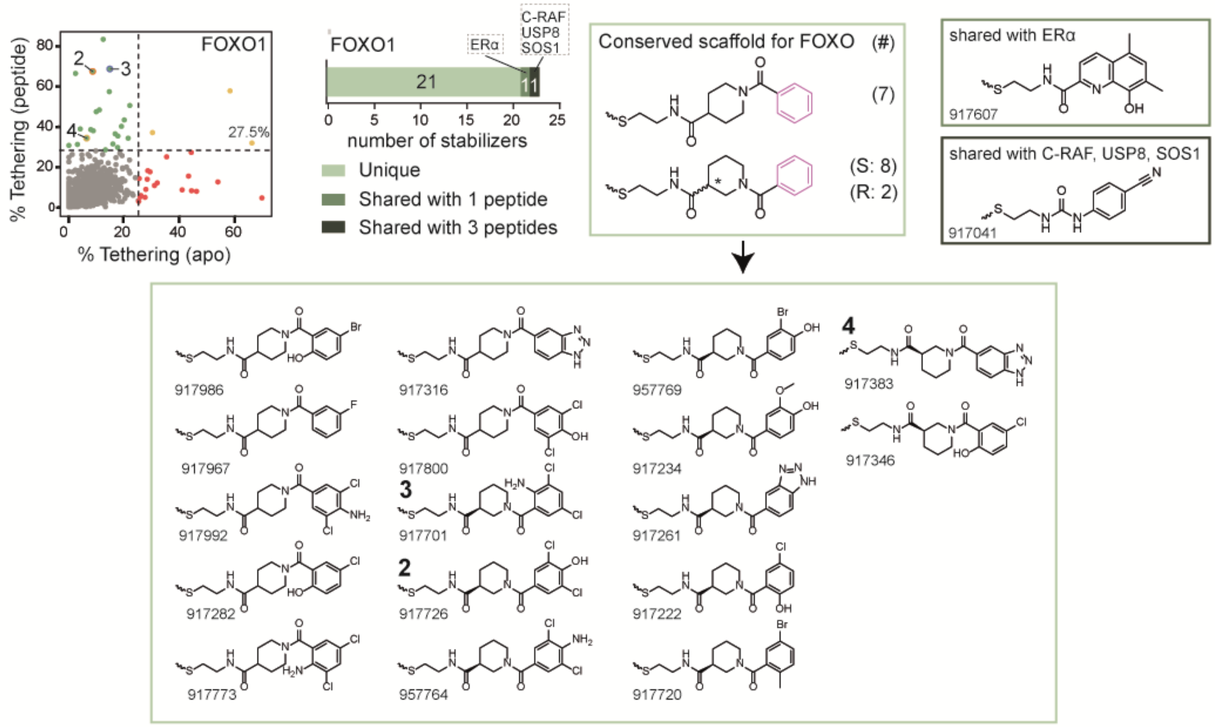


**Figure S6**: Scatter plot and chemical structures of the discovered stabilizers for FOXO1 from the primary screen, with in the light green box the unique stabilizers, the darker green box stabilizer shared with ERα and in the darkest green box the stabilizer shared with C-RAF, USP8 and SOS1.

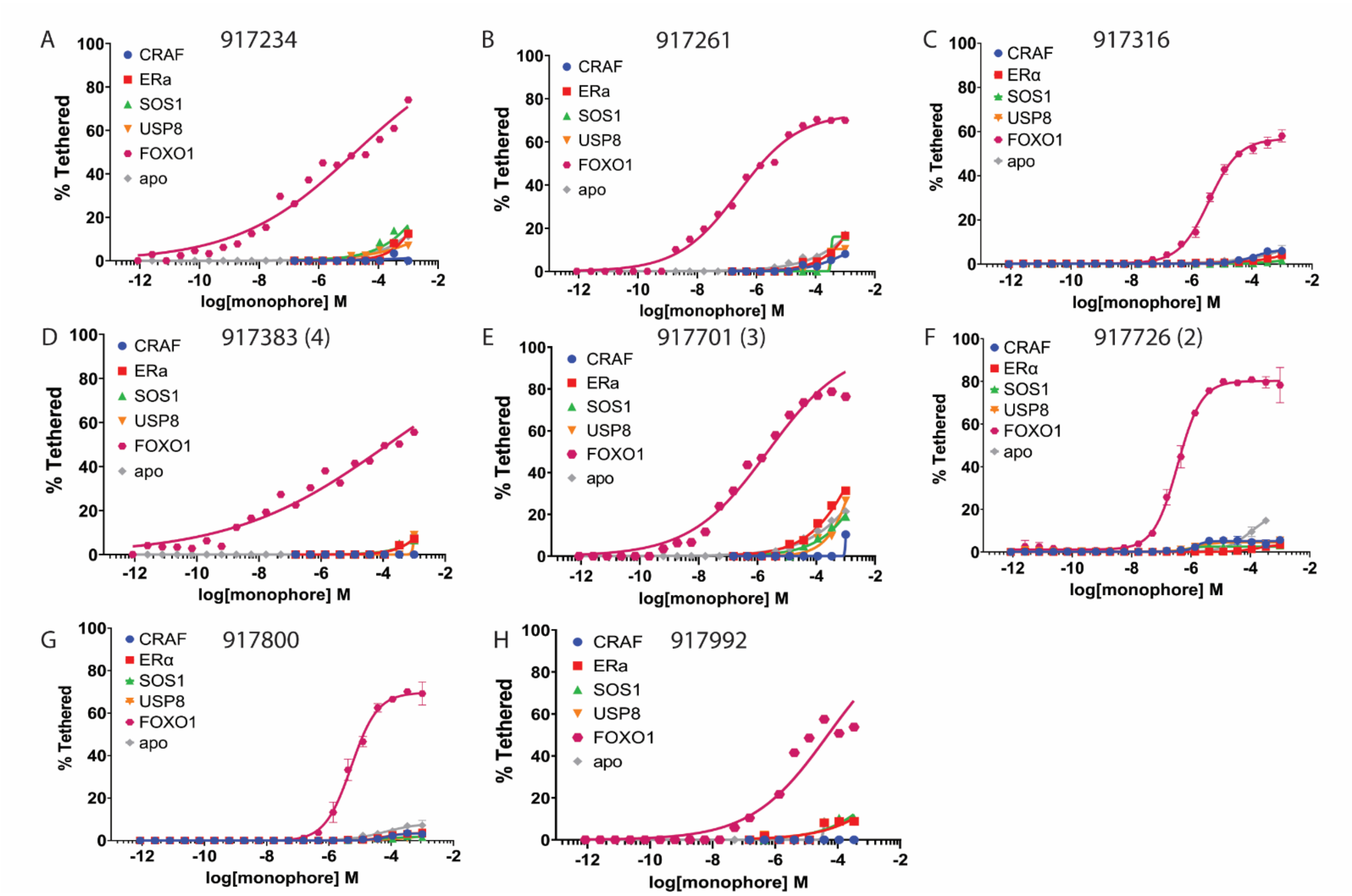


**Figure S7.** Mass spectrometry dose response (MSDR) curves (pink) for top FOXO1 selective stabilizers. (A) MSDR for FOXO1 hit compound 917234 showing selectivity for FOXO1 over other four 14-3-3σ client phospho-peptides and *apo*. $DR_{50}$: 16.9 μM. (B) MSDR for FOXO1 selective hit compound 917261. $DR_{50}$: 3.61 μM. (C) MSDR for FOXO1 selective hit compound 917316. $DR_{50}$: 42 nM. (D) MSDR for FOXO1 selective hit compound 917383 (compound **4** in main text). $DR_{50}$: 143.4 μM. (E) MSDR for FOXO1 selective hit compound 917701 (compound **3** in main text). $DR_{50}$: 2.2 μM. (F) MSDR for FOXO1 selective hit compound 917726 (compound **2** in main text). $DR_{50}$: 360 nM. (G) MSDR for FOXO1 selective hit compound 917800. (H) MSDR for FOXO1 selective compound 917992. $DR_{50}$: 48.7 μM.

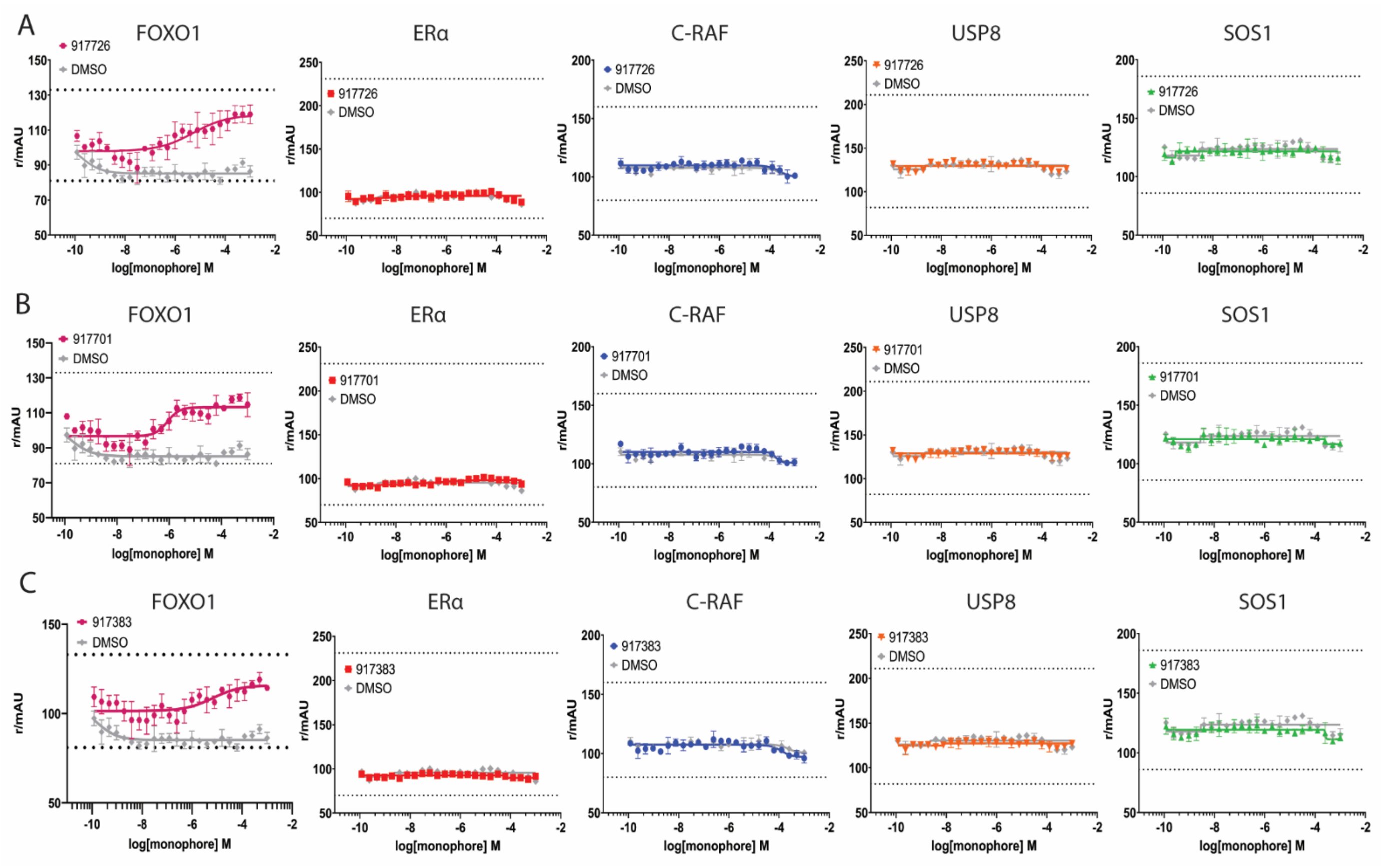


**Figure S8.** Fluorescence anisotropy dose response (FADR) curves for FOXO1 selective stabilizers 917726, 917701, and 917383 (compounds **2-4** in main text, respectively). Top black dashed line signifies maximum anisotropy control, the value of which was determined by peptide bound to 250 µM 14-3-3σ (at least 50-fold excess of peptide $K_D$). Bottom dashed line signifies minimum anisotropy control, value determined by peptide only and no 14-3-3σ. Controls were measured per plate. (A) 917726 dose-dependent stabilization of FOXO1 phospho-peptide (pink) binding to 14-3-3σ. $EC_{50}$ value with FOXO1: 5.1 µM. ERα (red), C-RAF (blue), USP8 (orange), and SOS1 (green) showed no increase in anisotropy over DMSO control (grey) indicating no stabilization. (B) 917701 stabilization of FOXO1 (pink) binding to 14-3-3σ. $EC_{50}$value with FOXO1: 850 nM. No increase stabilization over DMSO control (grey) observed for ERα (red), C-RAF (blue), USP8 (orange), or SOS1 (green). (C) 917383 stabilization of FOXO1 (pink) binding to 14-3-3σ. $EC_{50}$ value with FOXO1: 6.69µM. No increase stabilization over DMSO control (grey) observed for ERα (red), C-RAF (blue), USP8 (orange), or SOS1 (green).

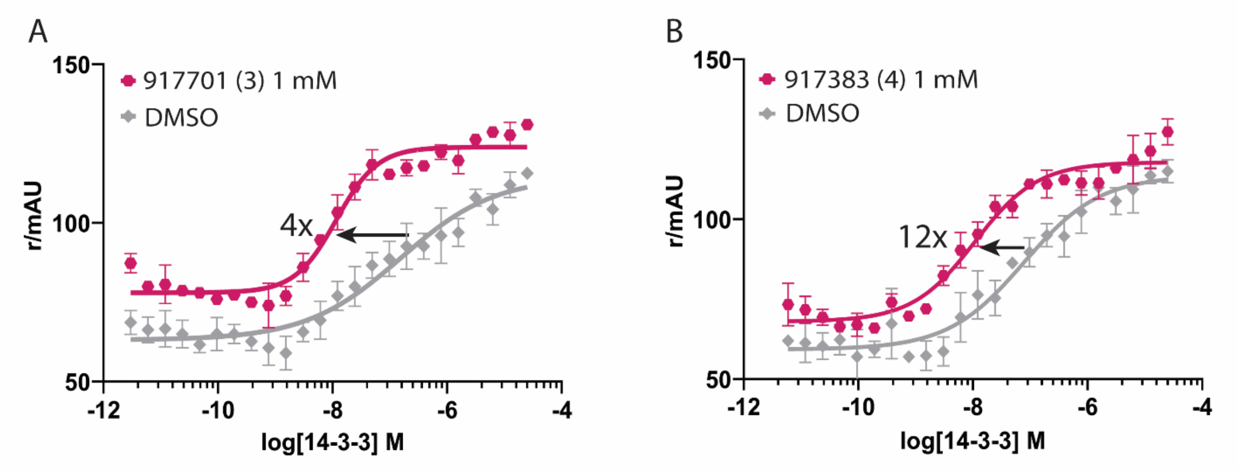


**Figure S9.** Fluorescence anisotropy protein titrations for FOXO1 and C-RAF selective stabilizers 917701 and 917383 (compounds **2** and **3** in main text, respectively. (A) Addition of 1 mM 917701 (compound **3**; pink curve) resulted in 4-fold decrease in 14-3-3σ/FOXO1 $K_D$ (10.6 nM vs. 42 nM) compared to DMSO control (grey curve). (B) Addition of 1 mM 917383 (compound **4**; pink curve) resulted in 12-fold decrease in 14-3-3σ/FOXO1 $K_D$ (9.7 nM vs. 111 nM)compared to DMSO control (grey curve).

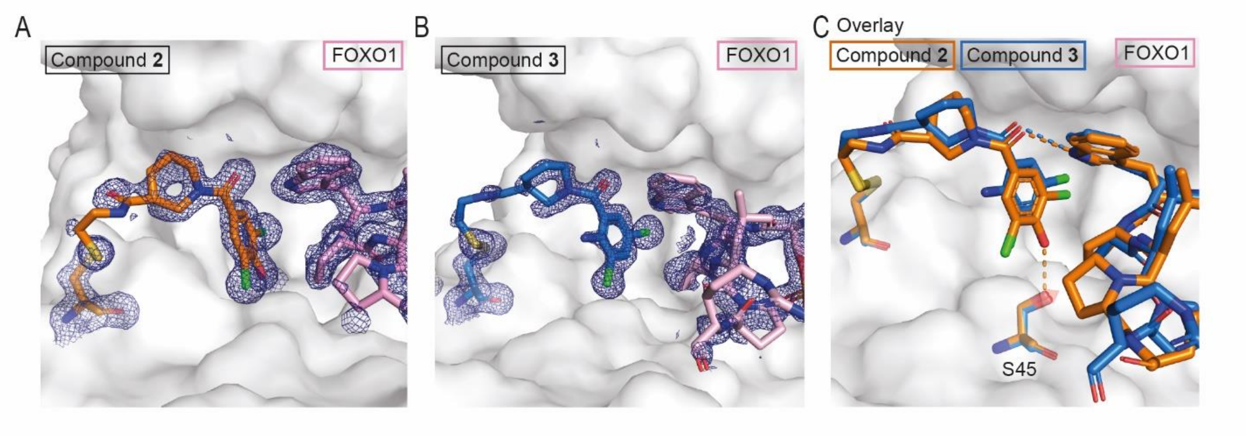


**Figure S10:** Crystal structures of (A) compound **2** (orange sticks) in complex with FOXO1 (pink sticks) and 14-3-3 (white surface) and (B) compound **3** (blue sticks) with FOXO1 (pink sticks) and 14-3-3 (white surface). (C) Crystallographic overlay of compound **2** (orange sticks) and compound **3** (blue sticks) with the FOXO1 peptide (peptide color matching compound complexed with) in 14-3-3 (white surface). Hydrogen interactions are depicted as orange dashed lines for compound **2** and blue dashed lines for compound **3**. 2Fo-Fc electron density maps are contoured at 1σ.

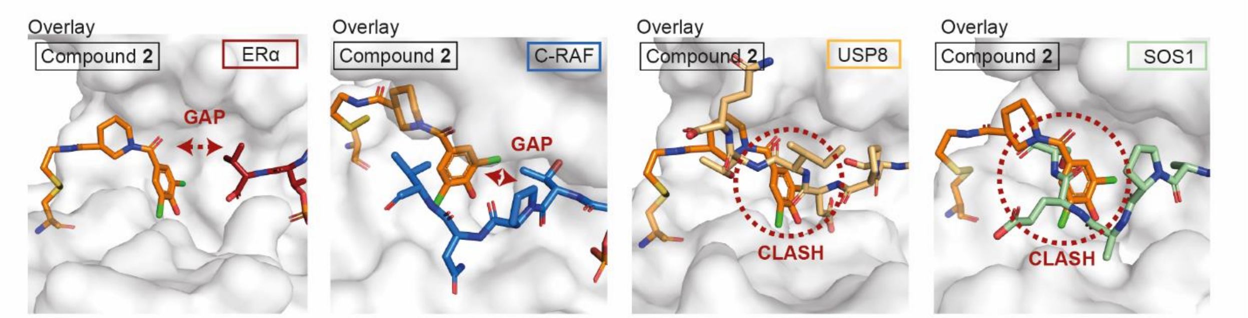


**Figure S11:** crystallographic overlay of compound **2** (orange sticks) with ERα (red sticks), C-RAF (blue sticks), USP8 (light orange sticks) and SOS1 (green sticks) in 14-3-3 (white surface). (PDB ID: 4JC3 (ERa), 4FJ3 (C-RAF), 6F09 (USP8), 6Y44 (SOS1).

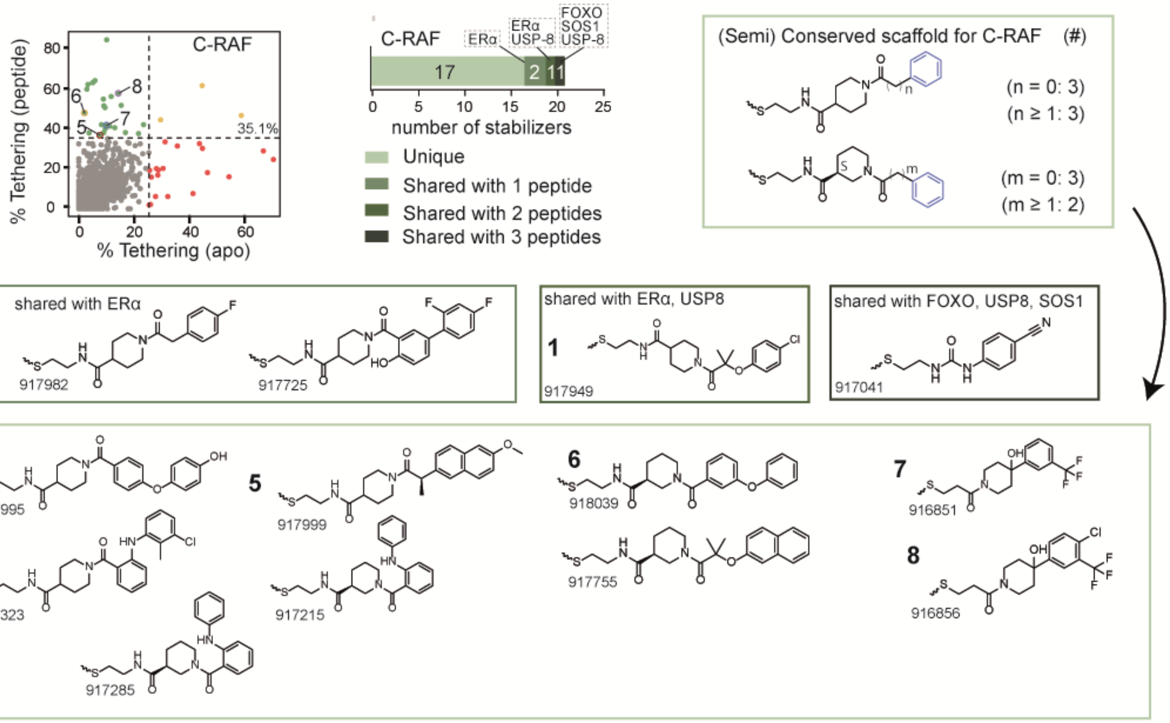


**Figure S12:** Scatter plot and chemical structures of the discovered stabilizers for C-RAF from the primary screen, with in the light green box the unique stabilizers, in darker green box the stabilizers shared with ERα, in the darker green box shared with ERα and USP8 (= compound **1**) and in the darkest green box shared with FOXO1, USP8 and SOS1.

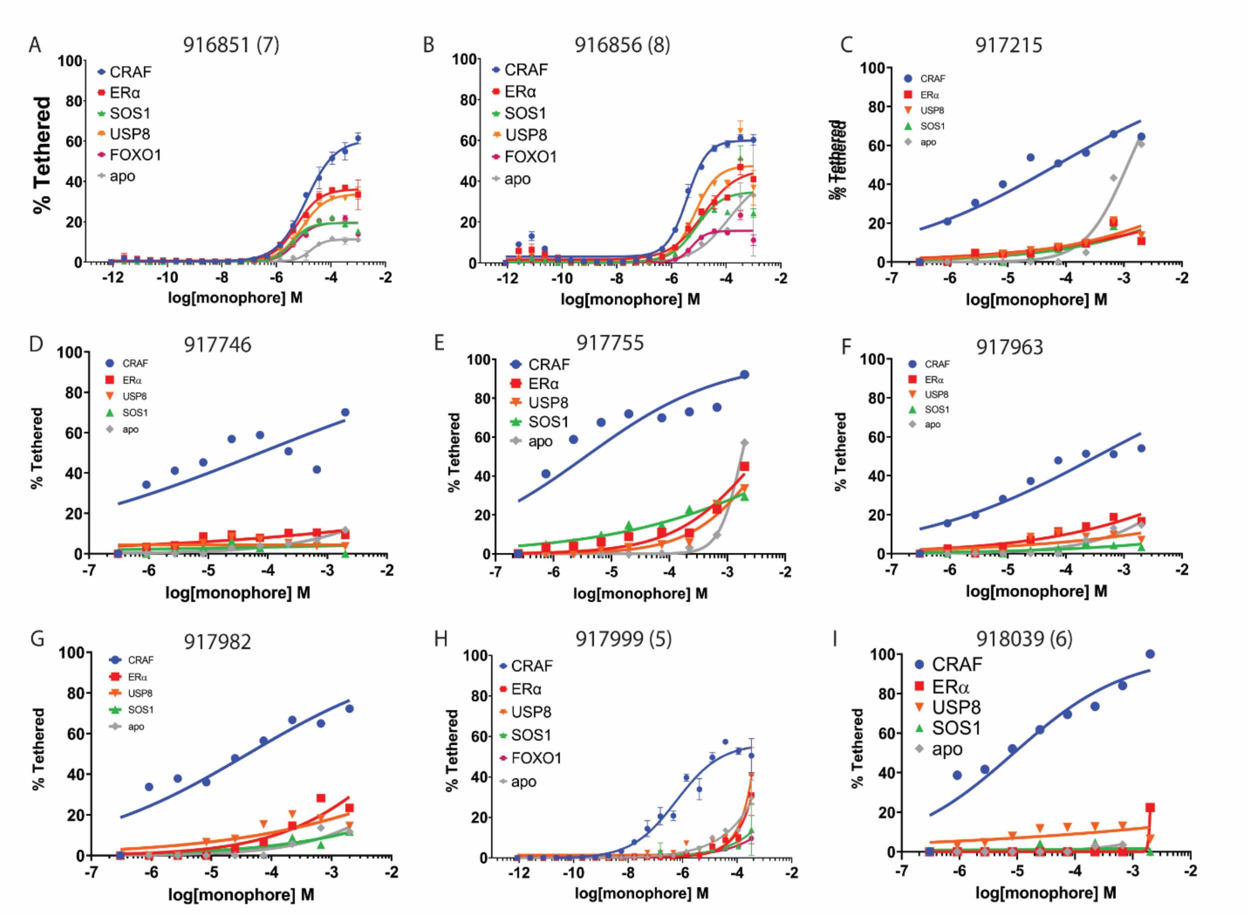


**Figure S13.** Mass spectrometry dose response (MSDR) curves (blue) for top C-RAF selective stabilizers. (A) MSDR for C-RAF hit compound 916851 (compound **7** in main text) showing preferential binding in presence of C-RAF phospho-peptide over other four 14-3-3σ client phospho-peptides and *apo*. $DR_{50}$ values: 12.2 μM (C-RAF), 5.62 μM (ERα), 8.54 μM (USP8), >2000 μM (SOS1), >2000 μM (SOS1), >2000 μM (apo). (B) MSDR for C-RAF hit compound 916856 (compound **8** in main text). 916856 was also a hit stabilizer in ERα primary screen and shows dose dependent engagement of 14-3-3σ in MSDR. $DR_{50}$ values: 4.96 μM (C-RAF), 14.5 μM (ERα), 7.05 μM (USP8), >2000 μM (SOS1), >2000 μM (FOXO1), >2000 μM (apo). (C) MSDR for C-RAF selective hit compound 917215. $DR_{50}$: 5.1 μM. (D) MSDR for C-RAF selective compound 917746. $DR_{50}$: 910 nM. (E) MSDR for C-RAF selective compound 917755. $DR_{50}$: 5.82 μM. (F) MSDR for C-RAF selective compound 917963. $DR_{50}$: 5.29 μM. (G) MSDR for C-RAF selective compound 917982. $DR_{50}$: 5.45 μM. (H) MSDR for C-RAF selective hit compound 917999 (compound **5** in main text). $DR_{50}$: 580 nM (I) MSDR for C-RAF selective hit compound 918039 (compound **6** in main text). $DR_{50}$: 8.78 μM.

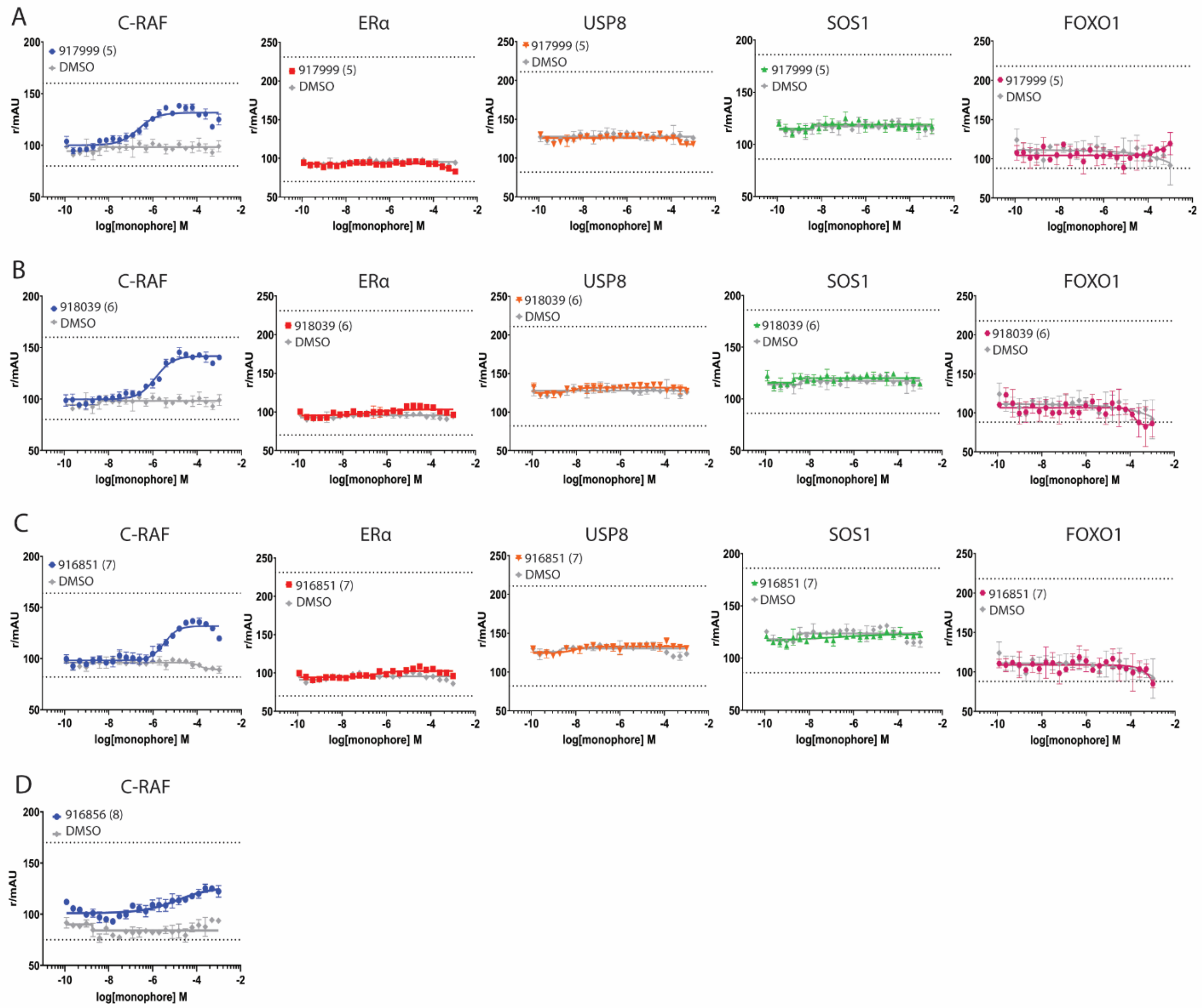


**Figure S14.** Fluorescence anisotropy dose response (FADR) curves for C-RAF selective stabilizers 917999, 918039, 916851, and 916856 (compounds **5-8** in main text, respectively). Top black dashed line signifies maximum anisotropy control, the value of which was determined by peptide bound to 250 μM 14-3-3σ (at least 50-fold excess of peptide $K_D$). Bottom dashed line signifies minimum anisotropy control, value determined by peptide only and no 14-3-3σ. Controls were measured per plate. (A) 917999 dose-dependent stabilization of C-RAF phospho-peptide (blue) binding to 14-3-3σ. $EC_{50}$ values with C-RAF: 220 nM. ERα (red), USP8 (orange), SOS1 (green), and FOXO1 (pink) showed no increase in anisotropy over DMSO control (grey) indicating no stabilization. (B) 918039 stabilization of C-RAF (blue) binding to 14-3-3σ.$EC_{50}$ value with C-RAF: 1.33 μM. No increased stabilization over DMSO control (grey) observed for ERα (red), USP8 (orange), SOS1 (green), or FOXO1 (pink). (C) 916851 stabilization of C-RAF (blue) binding to 14-3-3σ. $EC_{50}$ value with C-RAF: 3.18 μM. No increased stabilization over DMSO control (grey) observed for ERα (red), USP8 (orange), SOS1 (green), or FOXO1 (pink). (D) 916856 stabilization of C-RAF (blue) binding to 14-3-3σ. $EC_{50}$ value: 13.5 μM. Not sufficient compound for FADR with other peptides.

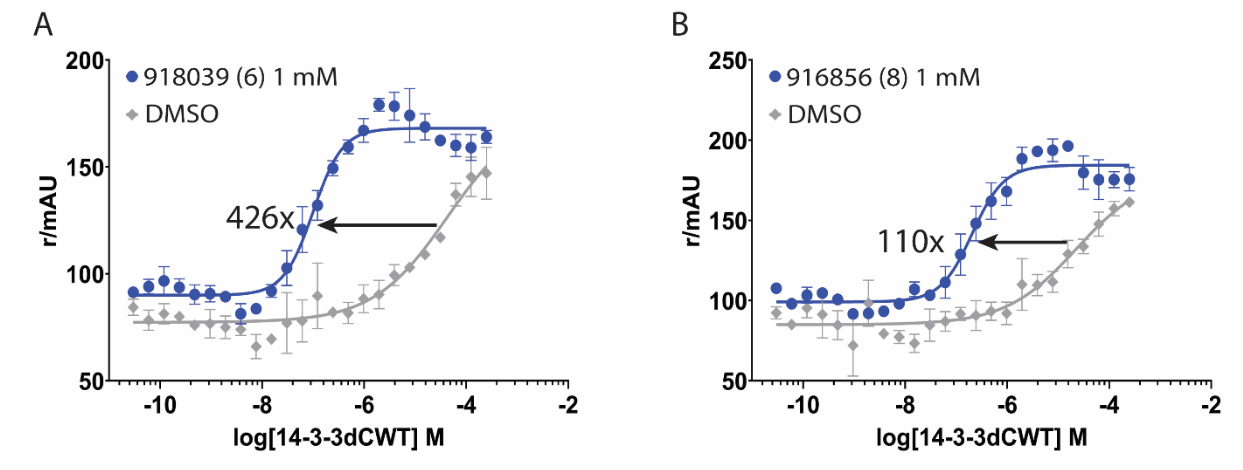


**Figure S15.** Fluorescence anisotropy protein titrations for FOXO1 and C-RAF selective stabilizers 918039 and 916856 (compounds **6** and **8** in main text, respectively). (A) Addition of 1 mM 918039 (compound **6**; blue curve) resulted in 426-fold decrease in 14-3-3σ/C-RAF $K_D$ (100 nM vs. 42 μM) compared to DMSO control (grey curve). (B) Addition of 1 mM 916856 (compound **8**; blue curve) resulted in 110-fold decrease in 14-3-3σ/C-RAF $K_D$ (207 nM vs. 23 μM) compared to DMSO control (grey).

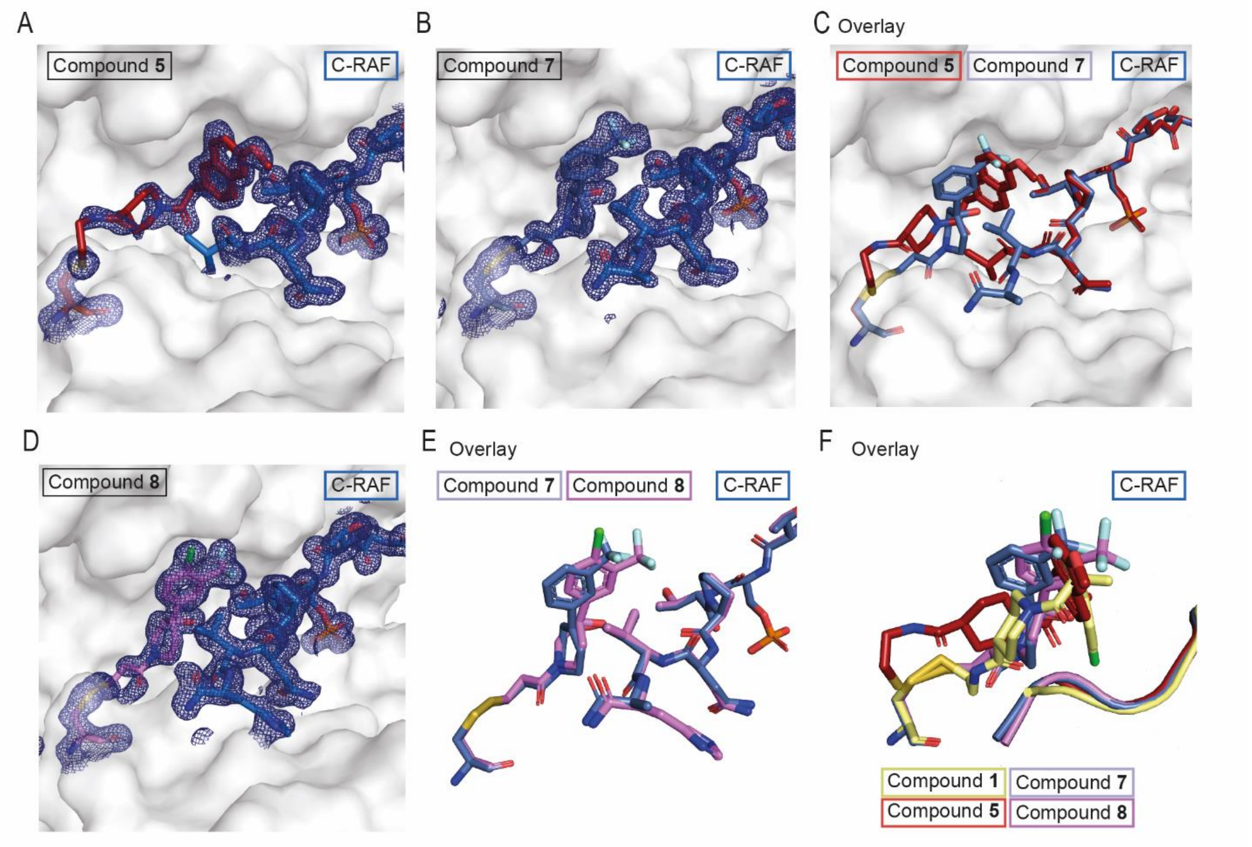


**Figure S16:** Crystal structures of stabilizers of C-RAF. (A) Electron density of compound **5** (red sticks) complexed with C-RAF (blue sticks) in 14-3-3 (white surface). (B) Electron density of compound **7** (light blue sticks) complexed with C-RAF (blue sticks) in 14-3-3 (white surface). (C) Crystallographic overlay of compound **5** (red sticks) and compound **7** (light blue sticks) with C-RAF (color peptide matching the compound complexed with) in 14-3-3 (white surface). (D) Electron density of compound **8** (violet sticks) with C-RAF (blue sticks) in 14-3-3 (white surface). (E) Crystallographic overlay of compound **7** (light blue sticks) and compound **8** (violet sticks) with C-RAF (color peptide matching the compound. (F) Crystallographic overlay of all crystallized stabilizers with C-RAF with compound **1** (yellow sticks), compound **5** (red sticks), compound **7** (light blue sticks) and compound **8** (violet sticks) with C-RAF represented as cartoon (color matching the compound complexed with) in 14-3-3 (white surface). 2Fo-Fc electron density maps are contoured at 1σ.

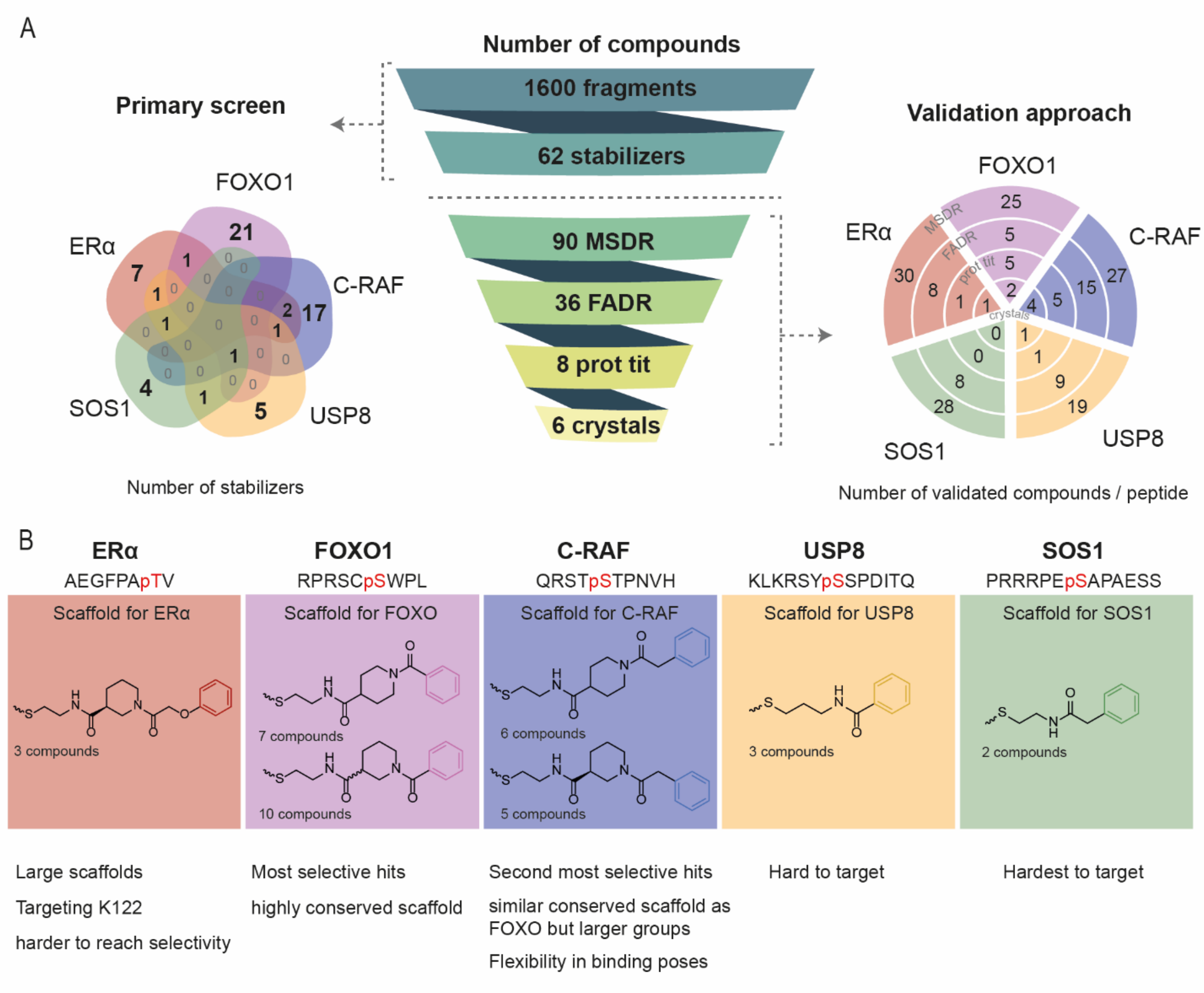


**Figure S17**: Overview of the screen performed with in (A) The screening funnel starting with the primary screen at the left depicted as a Venn diagram with the number of stabilizers discovered for each peptide and at the right the validation approach taken after the primary screen with the number of validated compounds per peptide per assay. (B) Conserved scaffold needed for stabilization of each peptide discovered from the primary screen.

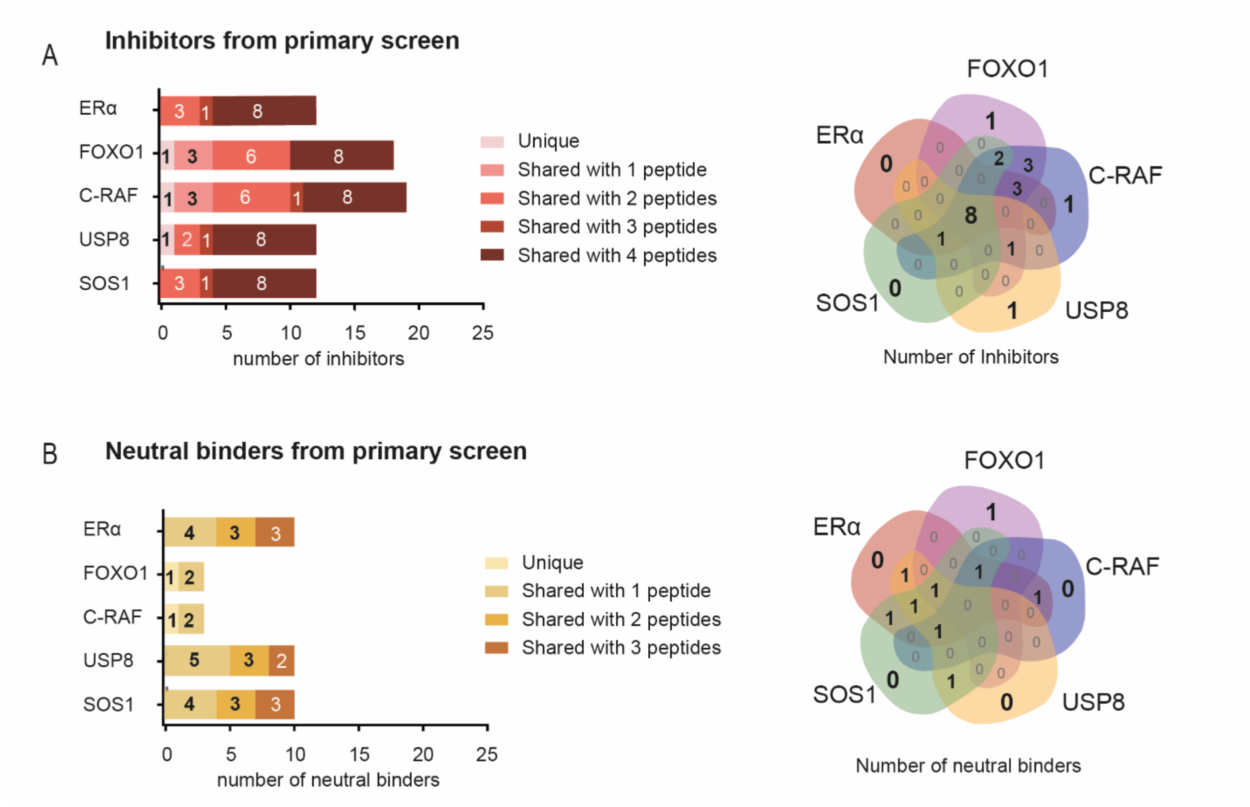


**Figure S18:** Overview of the number of discovered inhibitors and neutral binders from the primary screen. (A) Number of discovered inhibitors for each peptide depicted as (left) bar plots and (right) as a Venn diagram. (B) Number of discovered neutral binders for each peptide depicted as (left) bar plots and (right) as a Venn diagram.

Supporting References


(1) Sijbesma, E.; Hallenbeck, K. K.; Leysen, S.; de Vink, P. J.; Skóra, L.; Jahnke, W.; Brunsveld, L.; Arkin, M. R.; Ottmann, C. Site-Directed Fragment-Based Screening for the Discovery of Protein–Protein Interaction Stabilizers. *J. Am. Chem. Soc.* **2019**, *141* (8), 3524–3531. https://doi.org/10.1021/jacs.8b11658.

(2) Burlingame, M. A.; Tom, C. T. M. B.; Renslo, A. R. Simple One-Pot Synthesis of Disulfide Fragments for Use in Disulfide-Exchange Screening. *ACS Comb. Sci.* **2011**, *13* (3), 205–208. https://doi.org/10.1021/co200038g.

(3) Kabsch, W. XDS. *Acta Crystallogr. D Biol. Crystallogr.* **2010**, *66* (Pt 2), 125–132. https://doi.org/10.1107/S0907444909047337.

(4) Winn, M. D.; Ballard, C. C.; Cowtan, K. D.; Dodson, E. J.; Emsley, P.; Evans, P. R.; Keegan, R. M.; Krissinel, E. B.; Leslie, A. G. W.; McCoy, A.; McNicholas, S. J.; Murshudov, G. N.; Pannu, N. S.; Potterton, E. A.; Powell, H. R.; Read, R. J.; Vagin, A.; Wilson, K. S. Overview of the CCP4 Suite and Current Developments. *Acta Crystallogr. D Biol. Crystallogr.* **2011**, *67* (Pt 4), 235–242. https://doi.org/10.1107/S0907444910045749.

(5) Evans, P. R.; Murshudov, G. N. How Good Are My Data and What Is the Resolution? *Acta Crystallogr. D Biol. Crystallogr.* **2013**, *69* (Pt 7), 1204–1214. https://doi.org/10.1107/S0907444913000061.

(6) Vagin, A.; Teplyakov, A. MOLREP: An Automated Program for Molecular Replacement. *J. Appl. Crystallogr.* **1997**, *30* (6), 1022–1025. https://doi.org/10.1107/S0021889897006766.

(7) Emsley, P.; Cowtan, K. Coot: Model-Building Tools for Molecular Graphics. *Acta Crystallogr. D Biol. Crystallogr.* **2004**, *60* (Pt 12 Pt 1), 2126–2132. https://doi.org/10.1107/S0907444904019158.

(8) Moriarty, N. W.; Grosse-Kunstleve, R. W.; Adams, P. D. Electronic Ligand Builder and Optimization Workbench (ELBOW): A Tool for Ligand Coordinate and Restraint Generation. *Acta Crystallogr. D Biol. Crystallogr.* **2009**, *65* (Pt 10), 1074–1080. https://doi.org/10.1107/S0907444909029436.

(9) Adams, P. D.; Afonine, P. V.; Bunkóczi, G.; Chen, V. B.; Davis, I. W.; Echols, N.; Headd, J. J.; Hung, L.-W.; Kapral, G. J.; Grosse-Kunstleve, R. W.; McCoy, A. J.; Moriarty, N. W.; Oeffner, R.; Read, R. J.; Richardson, D. C.; Richardson, J. S.; Terwilliger, T. C.; Zwart, P. H. PHENIX: A Comprehensive Python-Based System for Macromolecular Structure Solution. *Acta Crystallogr. D Biol. Crystallogr.* **2010**, *66* (Pt 2), 213–221. https://doi.org/10.1107/S0907444909052925.

(10) Afonine, P. V.; Grosse-Kunstleve, R. W.; Echols, N.; Headd, J. J.; Moriarty, N. W.; Mustyakimov, M.; Terwilliger, T. C.; Urzhumtsev, A.; Zwart, P. H.; Adams, P. D. Towards Automated Crystallographic Structure Refinement with Phenix.Refine. *Acta Crystallogr. D Biol. Crystallogr.* **2012**, *68* (Pt 4), 352–367. https://doi.org/10.1107/S0907444912001308.

(11) Karplus, P. A.; Diederichs, K. Linking Crystallographic Model and Data Quality. *Science* **2012**, *336* (6084), 1030–1033. https://doi.org/10.1126/science.1218231.
